# Excess dietary sugar impairs colonic epithelial regeneration in response to damage

**DOI:** 10.1101/2021.08.18.456840

**Authors:** Ansen H.P. Burr, Junyi Ji, Kadir Ozler, Onur Eskiocak, Brian Yueh, Ashley V. Menk, Ana S.H. Costa, Natalie Rittenhouse, Chris W. Marshall, Pailin Chiaranunt, Lauren Mullinax, Abigail Overacre-Delgoffe, Vaughn S. Cooper, Amanda C. Poholek, Greg M. Delgoffe, Semir Beyaz, Timothy W. Hand

**Affiliations:** R.K. Mellon Institute for Pediatric Research, Pediatrics Department, Infectious Disease Section, UPMC Children’s Hospital of Pittsburgh, University of Pittsburgh, Pittsburgh PA, 15224; Department of Immunology, University of Pittsburgh, School of Medicine, Pittsburgh PA, 15261; School of Medicine, Tsinghua University, Beijing, 100084, China; Department of Microbiology, University of Pittsburgh, School of Medicine, Pittsburgh PA, 15261; Tumor Microenvironment Center, UPMC Hillman Cancer Center, Pittsburgh PA 15232; Cold Spring Harbor Laboratory, Cold Spring Harbor, NY, 11724

## Abstract

The colonic epithelium requires continuous renewal by intestinal stem cells (ISCs) to restore the barrier after damage and proliferation of epithelial cells is modulated by dietary metabolites. We demonstrate that mice fed a high sugar diet failed to repair colonic barrier damage, resulting in increased intestinal pathology. Culturing ISCs in excess sugar limited murine and human colonoid development, indicating that dietary sugar can directly affect colonic epithelial proliferation. Similarly, *in vivo* lineage tracing experiments and transcriptomic analysis indicated that dietary sugar impeded the proliferative potential of ISCs. ISCs and their immediate daughter cells predominantly rely on mitochondrial respiration for energy; however, metabolic analysis of colonic crypts revealed that a high sugar diet primed the epithelium for glycolysis without a commensurate increase in aerobic respiration. Colonoids cultured in high-glucose conditions accumulated glycolytic metabolites but not TCA cycle intermediates, indicating that the two metabolic pathways may not be coupled in proliferating intestinal epithelium. Accordingly, biochemically inducing pyruvate flux through the TCA cycle by inhibiting pyruvate dehydrogenase kinase rescued sugar-impaired colonoid development. Our results indicate that excess dietary sugar can directly inhibit epithelial proliferation in response to damage and may inform diets that better support the treatment of acute intestinal injury.

## Introduction

The modern diet of High-Income Countries is characterized by increased consumption of dietary fats and sugar, especially “acellular sugar” or simple carbohydrates that are readily absorbed by the host without additional digestive processing (Grundy et al., 2016; Monteiro, Moubarac, Cannon, Ng, & Popkin, 2013; Spreadbury, 2012). Indeed, the rate of sugar consumption has increased by 127% in the last 40 years, a trend that closely follows the rise in incidence of Inflammatory Bowel Disease (IBD) (Kearney, 2010). Diet is an important contributor to the development of IBD as epidemiological studies have found a positive association of IBD and high consumption of dietary sugar and sweetened beverages (Hou, Abraham, & El-Serag, 2011; Racine et al., 2016; Thornton, Emmett, & Heaton, 1979). Mice that consume a diet high in sugar have worse disease in models of colitis and clinical trials that significantly reduce dietary sugar have already shown promise in reducing disease burden in IBD patients in the pediatric intensive care unit (Laffin et al., 2019; Obih et al., 2016; Yeh et al., 2019). However, the mechanism behind this correlation remains unknown.

The intestinal barrier is exposed to billions of microorganisms, dietary products and their metabolites every day. To prevent barrier-failure and bacteremia, the intestinal epithelium is renewed every 3 to 5 days by the proliferative function of Lgr5^+^ intestinal stem cells (ISC) (N Barker et al., 2008; Nick Barker et al., 2007). ISCs reside at the base of crypts and asymmetrically divide to self-renew and to generate Transit Amplifying cells (TAs), which rapidly divide as they move up the crypt and differentiate into mature epithelial subsets such as goblet cells, enteroendocrine cells, and absorptive enterocytes (Bjerknes & Cheng, 1999; Cheng & Leblond, 1974). Rapid proliferation of crypts is particularly important after intestinal damage (Potten, 1990).

Diet-derived metabolites have been shown to directly alter both the proliferation and ‘stemness’ of ISCs. For example, calorie restriction leads to expansion of the ISC population and a reduction in other epithelial subsets, suggesting a preference for symmetric division rather than differentiation (Yilmaz et al., 2012). In contrast, a high fat diet increases both the self-renewal and proliferation capacity of ISCs due to a metabolic preference for fatty acid oxidation and aerobic respiration by TAs (Beyaz et al., 2016; Fan et al., 2015). The intestine is home to a large and diverse microbiota that aids in the digestion of food, in particular, the breakdown of fiber into short chain fatty acids (SCFA) which are also an important carbon source and fuel for fatty acid oxidation in the intestinal epithelium (Kelly et al., 2015; J. M. W. Wong, de Souza, Kendall, Emam, & Jenkins, 2006). Elucidating how our diet both directly and indirectly affects ISC function may have important implications for understanding how the intestine heals after damage caused by infection, Inflammatory Bowel Diseases (IBD) or after radiation therapy.

We show in a mouse model of intestinal damage (dextran sulfate sodium, DSS) (Chassaing, Aitken, Malleshappa, & Vijay-Kumar, 2014) that a high sugar diet induces worse colonic disease when compared to a high fiber diet and that excess sugar directly impairs the growth of intestinal stem cells cultured *in vitro*. Transcriptome and imaging data confirmed that high sugar diet inhibits the proliferation of ISCs and their crypt-resident daughter cells and that this phenotype is exacerbated by DSS-induced damage. Metabolic analysis of intestinal crypt cells showed that a HS diet skewed them towards glycolysis without a requisite ‘coupled’ increase in aerobic respiration. Indeed, forcing a coupling of glycolysis and the TCA cycle with the pyruvate dehydrogenase kinase inhibitor, DCA, restored the proliferative function of colonoids growing under high sugar concentrations. Together, these studies elucidate the potentially damaging effects a high sugar diet may have on the regenerative capacity of the intestinal epithelium after injury.

## Results

### High sugar diet leads to lethal colonic damage when treated with DSS

To determine the effect of excess dietary sugar on a murine model of intestinal damage, we fed C57BL/6 mice one of two defined diets: 1) high sugar (HS, 68%kcal from sucrose) or 2) high fiber diet (HF, 68%kcal from high amylose cornstarch) where macronutrient levels are equalized and differ only in the predominant source of carbohydrates (Table S1). Mice were fed a defined diet for two weeks then exposed to 3% dextran sodium sulfate (DSS) drinking water for one week. Compared with HF-fed and standard diet-fed (Std, chow in facility that is similar in composition to HF, see Table S1) controls, mice fed a HS diet had significantly greater weight loss and nearly 100% mortality by day 7 of DSS administration (Fig. 1 *A* and *B*). As weight loss is not the only, nor the best, indicator of intestinal damage, histological analysis demonstrated that, compared to HF or Std diet-fed mice, DSS-treated, HS-fed mice exhibited massive immune infiltration and loss of crypt structure in the colonic lamina propria by day 6 of DSS treatment (Fig. 1 *C* and *D*). This level of damage is normally seen at day 8-10 of DSS treatment, suggesting excess dietary sugar is accelerating disease progression (Nunes et al., 2018). Importantly, diet alone did not significantly alter the weights of diet-treated mice nor the amount of water consumed by each group, eliminating the possibility that weight loss was due to insufficient nutrition or greater DSS consumption (Fig. S1 *A* and *B*).

**Fig. 1.**
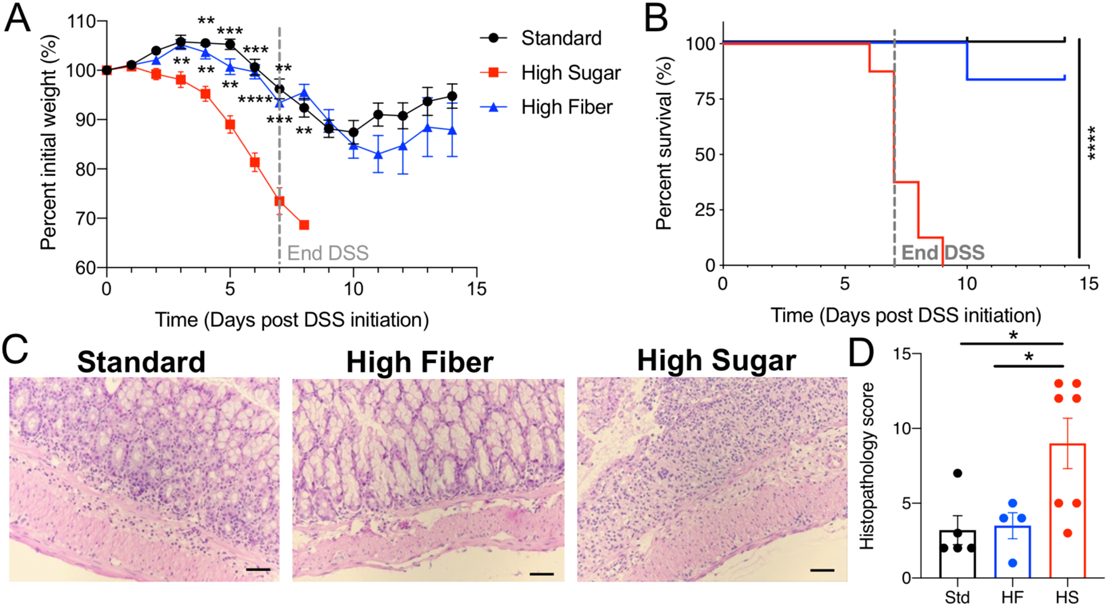
Excess dietary sucrose leads to lethal DSS-induced colonic damage. (*A-B*) 5-wk old female Taconic C57BL/6 mice were fed standard (Std), high fiber (HF), or high sugar diet (HS) for 2 weeks then treated with 3% DSS drinking water for 1 week to induce colonic inflammation. *(A*) Percent initial weight and (*B*) survival during and post DSS treatment. Data are representative of two independent experiments (n=4-5). Data points represent mean +/− SEM. Multiple t-tests performed against HS per day where *\*\*P<*0.01, *\*\*\*P<*0.001, *\*\*\*\*P<*0.00001. (C) Representative Hematoxylin and Eosin staining of colonic sections taken on day 6 of DSS treatment. Images were taken at 20X magnification (scale bar: 50μm). (D) Histopathology score of blinded H&E sections, where scores of 1-5 are mild colitis, 6-10 are moderate and 11-17 are severe, data are representative of 3 experiments (n=2-3). Data points represent mean +/− SEM. One-way ANOVA used to determine significance in (D) where *\*P<*0.05.

Sucrose is composed of two monosaccharides, glucose and fructose, which are differentially metabolized and absorbed in the intestine. We observed similar weight loss and lethal disease in mice drinking water supplemented with fructose, glucose or sucrose (10% by mass) when treated with DSS (Fig. S1*C*). Glucose appeared to have the most severe effects, with greater weight loss and mortality (Fig. S1*C*). However, all mice given sweetened water succumbed to DSS, while mice given unsweetened water lost less weight and recovered from colonic damage (Fig. S1*D*). However, mice prefer sweetened water (Sclafani, Zukerman, & Ackroff, 2014), and thus drink more sugar-sweetened water, which confounds interpretation of these experiments (Fig. S1*B*). Therefore, we focused on using HS and HF diets, where we have not observed differential water uptake and we can directly observe the effect of excess dietary sugar on the colonic response to damage (Fig. S1*B*).

### High sugar diet does not significantly alter the composition of the intestinal microbiome

We postulated that sugar may be altering the intestinal microbiota of HS-fed mice by expanding the population of *Enterobacteriaceae*, which can thrive on simple carbohydrates (Ayres, Trinidad, & Vance, 2012; Kamada, Chen, Inohara, & Núñez, 2013). Although 16S rRNA-sequencing of fecal samples showed that defined diets altered the intestinal microbiota compared to mice fed the standard diet (Std), there was no significant difference between the microbiota of HS− or HF− fed mice after two weeks of consuming their defined diets as determined by measurement of diversity, Principal Coordinate and LEfSe analysis (Fig. 2 *A-C* and data not shown). Seven days after inflammation was introduced with DSS, we observed an outgrowth of *Enterobacteraceae* and *Enterococcaceae* in the microbiota of HS-fed mice (Fig. 2*D*). However, these same bacterial taxa are expanded under a variety of inflammatory intestinal conditions of varying severity, and thus are unlikely to be the sole cause of the rapid failure of the colonic epithelium seen in DSS-treated HS-fed mice (Lupp et al., 2007; Winter et al., 2013). Further, mice receiving sugar in their water bottle succumbed to DSS treatment but did not exhibit the same outgrowth of *Enterobacteraceae* and *Enterococcaceae*, indicating that it is not necessary for lethal sequelae (Fig. 2*D*). To test whether shifts in the microbiome of HS-fed mice exacerbated DSS-colitis, we transferred the fecal microbiome from HS-fed mice and from HF-fed mice into germ-free mice receiving standard chow and treated each group with DSS. Mice receiving the HS microbiota showed decreased survival compared to germ-free mice that received HF microbiota (Fig. 2 *E-F*). However, mice that received the microbiota from HS-fed mice did not lose weight as quickly as HS-fed mice and had increased length of survival, suggesting that the microbiota is not sufficient to induce the acute negative effects of excess dietary sugar. In previous studies, both a fiber-free diet and drinking water supplemented with sugar contributed to the expansion of mucus degrading bacteria and increased their susceptibility to colonic bacterial infection (Desai et al., 2016; Khan et al., 2020). In our experiments, despite HS diet containing low levels of fiber, the frequency of *Akkermansia spp.* did not discriminate HS or HF-fed mice and thus could not explain the phenotype of DSS-treated HS-fed mice (Fig. 2*G*).

**Fig. 2.**
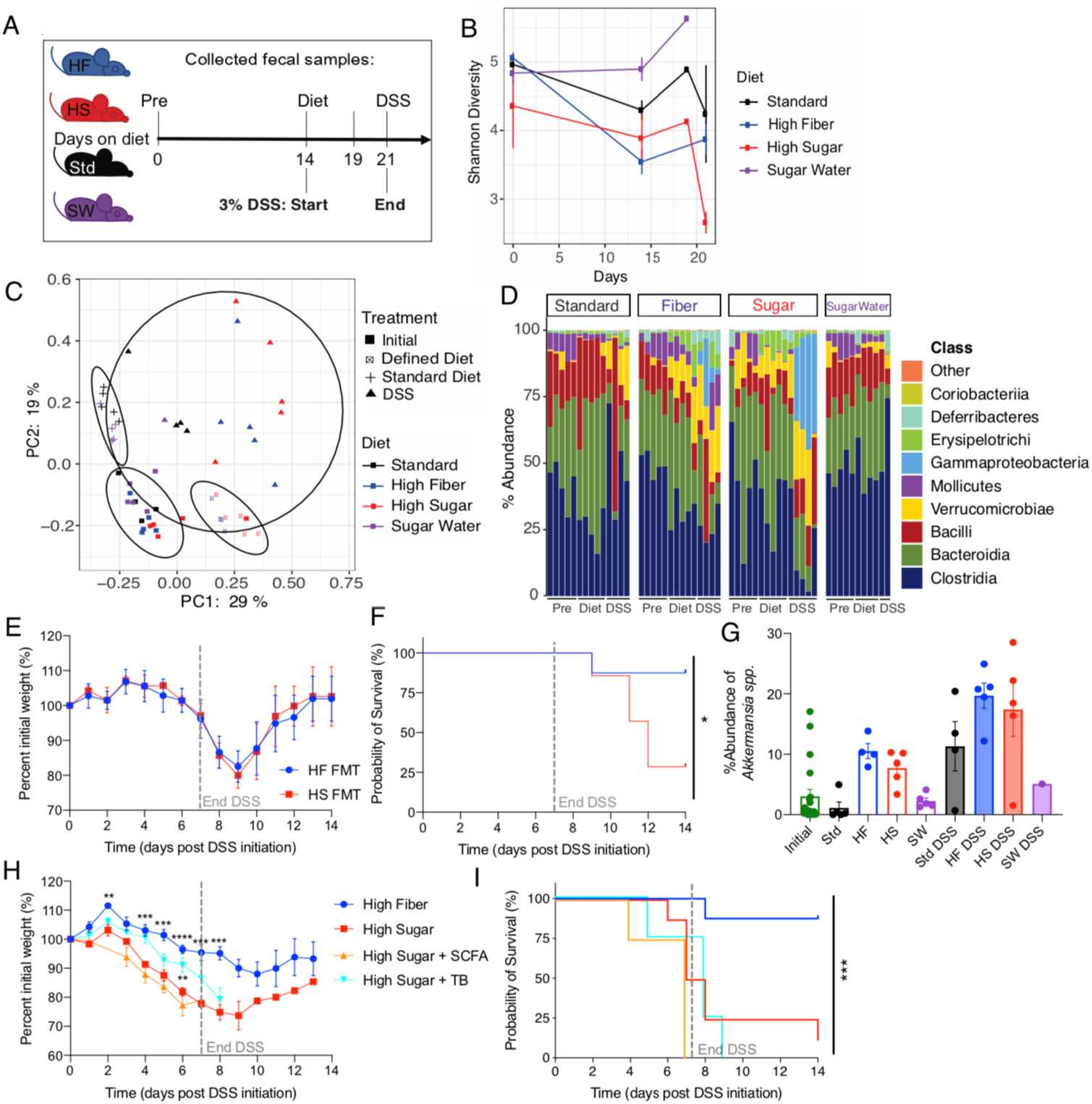
High sugar diet-associated shifts in the microbiome are not sufficient to induce the rapid lethal colitis phenotype of HS-fed, DSS-treated mice. (*A*-*D,G*) 5-wk old female Taconic C57BL/6 mice were fed HS or HF diet for 2 weeks then treated with 3% DSS drinking water, fecal samples were collected for 16S sequencing throughout. (*A*) Schematic of diet and DSS treatment and days fecal samples were collected for 16S rRNA analysis. “Pre” refers to sample collection on day mice arrived at our facility, “Diet” refers to 14 days of the respective diet (HF=high fiber, HS= high sugar, Std=standard facility chow, SW=Std with 10% sucrose in water) and “DSS” refers to samples collected during DSS treatment. (*B*) Shannon Diversity of microbial community over time, data points represent mean +/− SD. (*C*) Ordination plot based on the Principle Coordinate Analysis (PCoA) (Bray Curtis) demonstrate taxonomic variations of microbial communities across mice of different diet treatments where “Defined Diets” include HF and HS, and “Standard Diet” refers to Std and SW. (*D*) Relative abundances of top 10 most abundant bacterial classes. (*G*) Relative abundance of *Akkermansia spp.* as determined by 16S rRNA gene sequencing. (*E-F*) Mice were fed HF, HS, or HS diet with short chain fatty acid (SCFA) supplementation in the water or tributyrin (TB) supplemented in the diet for 1 week then treated with 3% DSS drinking water (dotted line) for 1 week to induce colonic damage. (*E*) Percent initial weight and (*F*) survival curve over DSS treatment duration are shown. (H-I) Germ-free female C57BL/6 mice were gavaged with fecal microbiome from HS or HF-fed SPF C57BL/6 mice, then treated with 3% DSS while fed Std. diet. (*H*) Representative weight loss curve and (*I*) survival are shown. Data are representative of two independent experiments (n=4). Data points represent mean +/− SEM. Multiple t-tests performed against HS per day where *\*P<*0.05, *\*\*P<*0.01, *\*\*\*P<*0.001, *\*\*\*\*P<*0.0001.

### Short chain fatty acid supplementation cannot prevent lethal complications of DSS-colitis in sugar-fed mice

Short chain fatty acids (SCFA) are an important microbiome-derived byproduct of dietary fiber that are absorbed by the host to provide nutrients to colonocytes, support the expansion of T regulatory cells, and dampen inflammation derived from innate immune cells (Donohoe et al., 2011; Furusawa et al., 2013; Kelly et al., 2015; J. M. W. Wong et al., 2006). Given the HS diet has less fiber, it may provide fewer SCFAs to the host. However, the addition of SCFA in the water of HS-fed mice did not save them from lethal colitis (Fig. 2 *H* and *I*). To ensure that SCFA were targeted specifically to the colon, rather than getting absorbed entirely by the small intestine, we also supplemented the HS diet with tributyrin (TB), which is broken down into butyrate and absorbed in the colon (Byndloss et al., 2017; Kelly et al., 2015). Tributyrin supplementation also did not rescue the HS-fed mice as they exhibited the same weight loss and lethality as mice on HS diet alone (Fig. 2 *H* and *I*), suggesting that it is not the relative lack of fiber and SCFA byproducts that is detrimental to colonic health during intestinal damage in our model, but the excess sugar.

### Short-term high sugar diet does not increase blood sugar or intestinal permeability

Excess dietary sugar has been linked to systemic diseases, such as diabetes, and elevated blood glucose can lead to impaired intestinal integrity (Thaiss et al., 2018). However, after two weeks of consuming the defined diets, we detected no differences in the fasted or post-prandial blood glucose levels (Fig. S2 *A* and *B*) or in intestinal permeability of HS or HF-fed mice prior to DSS treatment (Fig. S2*C*). Therefore, high levels of blood glucose and altered intestinal permeability could not explain the increased susceptibility of HS-fed mice to intestinal damage.

### Dietary sugar must be present during colonic damage for lethal disease and can be absorbed by the colonic epithelium

We next hypothesized that if sugar was directly affecting the epithelium, then HS diet may need to coincide with DSS-induced intestinal damage. To test this, we fed mice Std or HS diet for 2 weeks then reversed the diets on the first day of DSS treatment. Mice that were fed HS diet for 2 weeks and switched to the Std diet with the initiation of DSS lost weight similar to the group fed Std diet throughout, while mice fed Std diet and switched to HS diet during DSS treatment lost weight similar to the group fed HS throughout the experiment (Fig. S2*D*). Therefore, excess sugar must be present contemporaneously to exacerbate DSS-induced disease. To directly test whether the colonic epithelium can absorb luminal glucose we utilized Lgr5 reporter mice (*Lgr5^eGFP-cre-ERT2^*) and fluorescently labelled glucose model (GlucoseCy5) (Watson et al., 2021). With this approach, we detected colonic epithelial uptake of fluorescent glucose introduced via enema that reached the Lgr5^+^ colonic stem cells, suggesting that crypt cells can directly uptake luminal glucose (Fig. S2*E*). The function of the colon is to remove water and electrolytes while the absorption of nutrients and metabolites is performed by the small intestine. These functional differences are demonstrated by the expression of glucose transporters, as the colonic epithelium only expresses glucose transporters that bring glucose into the cell from the lumen (*Slc5a1*; SGLT1) and from the blood (*Slc2a1*; GLUT1), whereas the small intestinal epithelium also expresses GLUT2 (*Slc2a2*), the bi-directional glucose transporter that exports glucose out of the cell and into the bloodstream (Fig. S2*F*) (Röder et al., 2014; Wang et al., 2020; Yoshikawa et al., 2011). Thus, the colonic epithelium is unable to shuttle glucose out of the cell by the canonical pathway and the colon is likely a final destination for absorbed sugar.

### Excess sugar impairs development of colonoids *in vitro*

To determine the direct impact of excess sugar on the function and development of colonic epithelium, we utilized murine colonoids, 3-dimensional epithelial structures that are generated from colonic crypts isolated from mice. We found that culturing crypts into colonoids in 50mM or more of sucrose, fructose or glucose led to fewer and smaller colonoids that had decreased viability compared to cells cultured in 12.5mM of sugar, in a dose-dependent manner (Fig. 3 *A* and *B*). Reduced colonoid growth was not due to osmotic pressure as fully developed colonoids were viable when cultured in greater than 100mM of sucrose, fructose or glucose (Fig. S3, *A-C*). Similar results were observed with human colonoids, where colonoids that had been dispersed and regrown had inhibited growth into robust colonoid structures when exposed to high sugar concentrations, while fully developed human colonoids had no change in viability or number (Fig. 3 *C* and *D* and Fig. S3 *D* and *E*). These results demonstrate a direct impairment of epithelial development and proliferation in high sugar conditions, both in mouse and human colonoids, that acts independently of the microbiome and local immune cells.

**Fig. 3.**
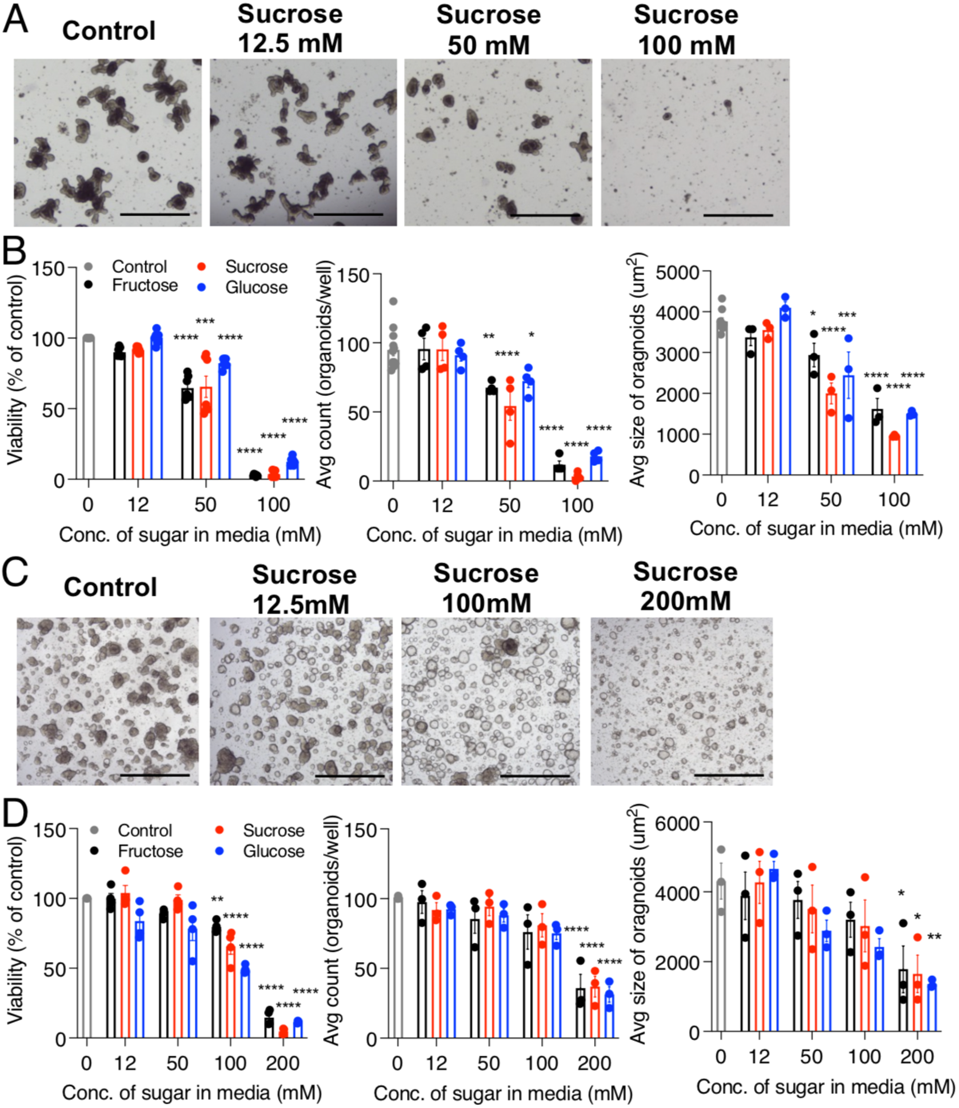
Excess sugar directly impairs *in vitro* colonoid formation by Lgr5^+^ intestinal stem cells. (A) Murine colonic crypts were cultured in increasing concentrations of sucrose, glucose or fructose under conditions that promote colonoid developed. Shown are representative images of colonoids 5 days after seeding. Images were taken at 4X magnification (scale bar= 200μm). (B) Viability (percent CTG luminescence of control), number and size of colonoids after 5 days in culture. (C) Patient colon samples were grown in culture conditions that promote colonoid growth and then dispersed to single cells and regrown with increasing concentration of sugar in the media for 12 days. (D) Average viability (percent CTG luminescence of control), number and size of colonoids after 12 days in culture are shown. Data are representative of 2 experiments (n=3) and data points are mean +/− SEM. Stats represent one-way ANOVA with multiple comparisons to Control, where \**P*<0.05, \*\**P*<0.01, \*\*\**P*<001, \*\*\*\**P*<0001.

### Excess dietary sugar impairs the epithelial proliferative response to damage

To confirm if HS diet similarly alters the regeneration of the colonic epithelium *in vivo*, we measured the transcriptome (RNAseq) of the colonic epithelium from *Rag1*^-/-^ mice (to ensure no intraepithelial lymphocyte contamination) fed HS or HF diet for 2 weeks. Critically, B and T cells are not required for the effects of HS diet as *Rag1^-/-^* mice phenocopy C57BL/6 mice after treatment with 3% DSS (Fig. S4 *A* and *B*). With diet alone, there were few transcriptional changes when comparing the intestinal epithelium of HS and HF-fed mice (Fig. 4*A* and Fig. S4 *C* and *D*). In contrast, after 3 days of DSS-induced damage, the transcriptome of the colonocytes from DSS-treated, HS-fed mice showed a reduction in the expression of the core gene signatures of Lgr5^+^ ISCs, TA cells, and secretory goblet cells, compared to DSS-treated HF-fed mice (Fig. 4*A* and Fig. S4 *E* and *F*). The genes associated with enteroendocrine cells were not substantially affected, indicating that the effect of HS diet is selective to specific epithelial cell types (Fig. 4*A*). These results were confirmed using Gene Set Enrichment Analysis (GSEA) which demonstrated an enrichment for Lgr5^+^ intestinal stem cell signature in DSS treated HF-fed epithelium compared to DSS-treated HS epithelium as well as enrichment in gene sets involved in cell cycle progression and proliferation, including E2F targets, G2M checkpoint genes, Myc targets, Cell Cycle genes, DNA repair and Mitotic spindle gene sets (Fig. 4*B* and Fig. S4*G*) (Muñoz et al., 2012). In contrast, epithelium from HS-fed DSS-treated mice showed an enrichment for the Epithelial-Mesenchymal transition gene set, which is a characteristic process of Crohn’s Disease, a subset of IBD, leading to fibrosis and stricturing (Fig. S4*G*). Typically, *Lgr5* expression and function is reduced by day 7 of DSS treatment, yet HS-fed mice exhibit a loss of *Lgr5* expression by day 3 of DSS treatment, indicating that dietary sugar accelerates disease progression in this model (Fig. 4 *A* and *B*) (Harnack et al., 2019). HS/DSS-treated mice also display reduced *Atoh1* expression which may indicate impaired production of the earliest secretory-progenitor cells from Lgr5^+^ ISCs in these mice (Fig. 4*A*). Proliferation is required to restore damaged epithelium, therefore we hypothesized that sugar may be impairing the regenerative response of the epithelium. Indeed, colonic epithelium from HS-fed mice expressed lower levels of the proliferative marker Ki67 after 3 days of DSS exposure and had fewer total epithelial cells by day 4 of DSS treatment, suggesting an impaired proliferative response when exposed to DSS-damage (Fig. 4 *C-E*). Additionally, we observed no differences in TUNEL and activated Caspase-3 staining after 3 days of DSS treatment, indicating that HS diet was not increasing cell death in the colonic epithelium (Fig. S5 *A-C*).

**Fig. 4.**
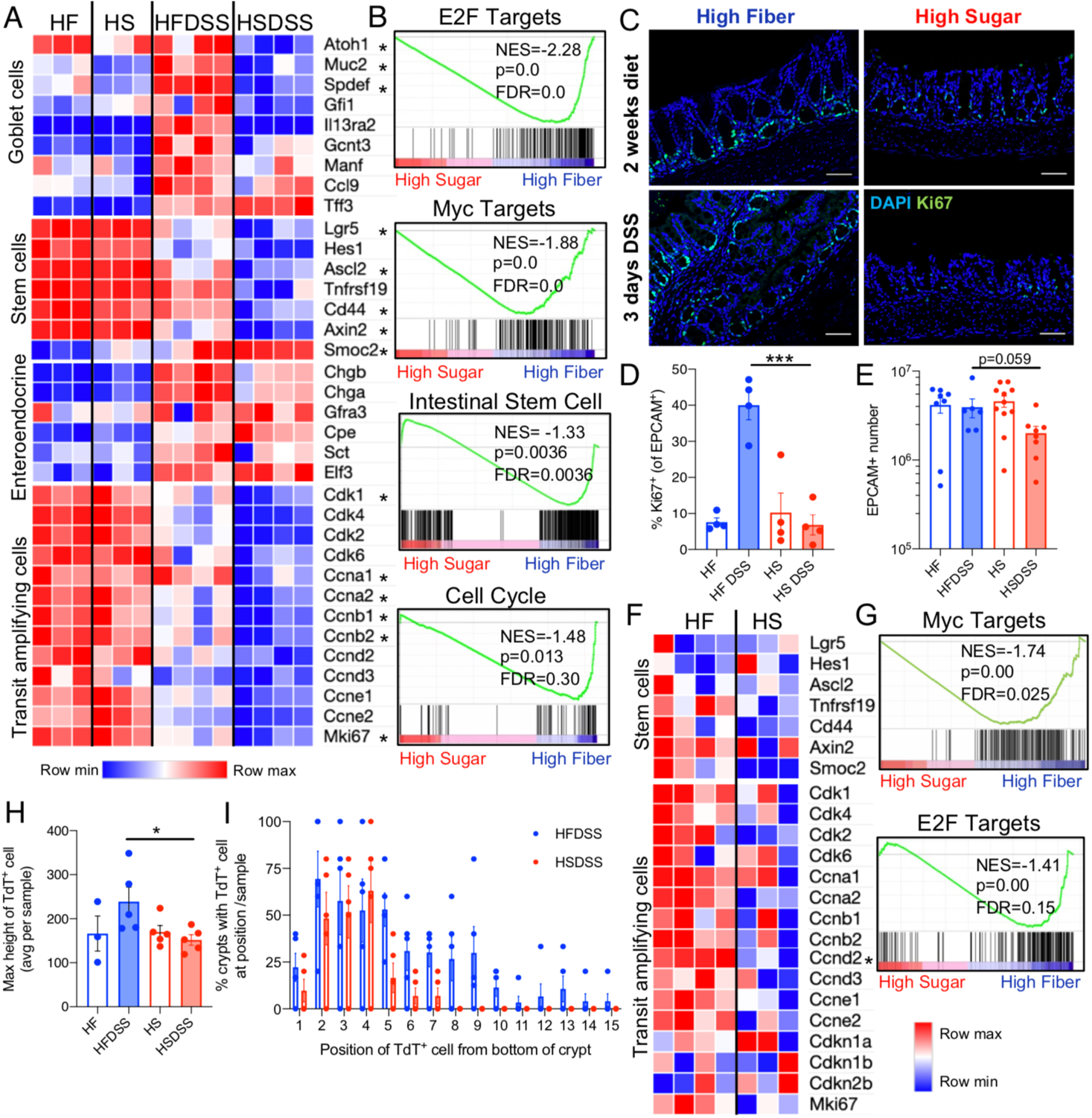
High sugar diet impairs the proliferation of intestinal stem cells. (*A-B*) Colonic epithelium was isolated from HS or HF-fed *Rag1*^-/-^ female mice with or without 3 days of 3% DSS treatment and analyzed by RNAseq(n=3-4). (*A*) Expression level of epithelial subset gene signatures from bulk colonic epithelial RNAseq, where red and blue represent high or low expression level, respectively, normalized across rows and * represents genes that are significantly differentially expressed in control versus glucose treated colonoids (FC>1.5, *P*<0.05, FDR<0.3). (*B*) Gene set enrichment analysis (GSEA) of colonic epithelium RNAseq data showing enrichment of genes in HF DSS treated or HS DSS treated mice for gene sets as indicated. (C) Representative images of colonic sections stained for Ki67 (green) from mice fed HS or HF diet for two weeks and treated for 3 days with 3% DSS. Images were taken at 20X magnification (scale bars=50μm). (D) Mice were fed HS or HF diet and treated 3 days with 3% DSS, colonic epithelium was isolated and stained with Ki67 for flow cytometric analysis. Data points represent percent of EPCAM^+^ cells that are Ki67^+^ per colonic sample, error bars are SEM, representative of 2 experiments (n=4). One-way ANOVA test was used to determine significance where \*\*\**P*<0.001. (E) Number of EPCAM^+^ cells in colon after 4 days of 3% DSS treatment. Data are representative of 2 experiments (n=3-5) and data points represent mean +/− SEM. Student’s t-test used to determine significance. (*F*-*G*) Lgr5^+^ intestinal stem cells (ISC) were isolated from Lgr5^eGFP-Cre-ERT2^ female mice fed HS or HF diet for 2 weeks and analyzed by RNAseq (n=3-4). (H) Transcript expression level of epithelial subset gene signatures as determined by Lgr5^+^ ISC RNAseq, where red and blue represent high or low expression level normalized across rows, respectively rows and represents genes that are significantly differentially expressed in control versus glucose treated colonoids (FC>1.5, *P*<0.05, FDR<0.3). (*G*) GSEA of Lgr5^+^ ISC RNAseq data showing enrichment of genes in HF-fed or HS-fed mice for gene sets as indicated. (*H-I*) *Lgr5^eGFP-Cre-ERT2^ Rosa^LSL-TdTomato^* mice were fed HS or HF diet for 2 weeks, injected with Tamoxifen to induce Tomato expression (*H*) the percent of GFP^+^ crypts containing Tomato^+^ progeny at the specified position along crypt and (*I*) height of most distant Tomato^+^ progeny from bottom of crypt (averaged per GFP^+^ crypt). Data are representative of two independent experiments (n=2-3) and data points represent mean +/− SEM. One-way ANOVA used to determine significance where *\*P<*0.05.

**Fig. 5.**
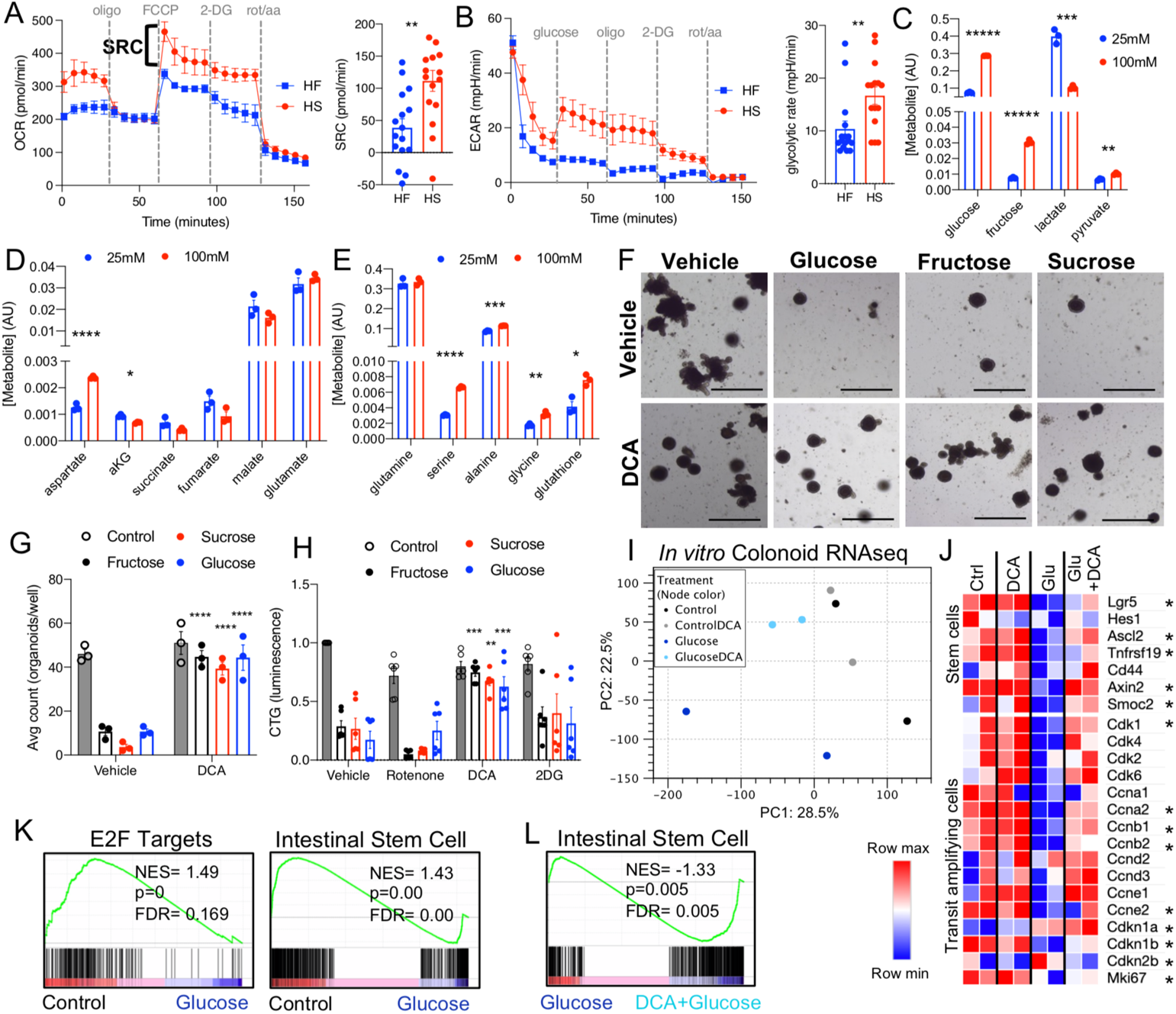
High sugar diet modulates the metabolic capacity of colonic crypts. (*A-B*) Colonic crypts were isolated from mice fed HS or HF diet for 2 weeks and plated on Matrigel coated Seahorse XF analyzer plate. (A) Representative oxygen consumption rate (OCR) trace and tabulated spare respiratory capacity (SRC: difference between basal and maximal oxidative rates, achieved after FCCP injection). *(B*) Representative extracellular acidification rate (ECAR) trace was measured after 3 hours of glucose deprivation. Glycolytic rate was measured by subtracting the basal rate after 2-DG injection from the maximum response post glucose injection. (*A-B*) For Seahorse traces, data are representative of four experiments (n=4) and data points represent mean +/− SEM and tabulated bar charts represent mean +/− SEM with each point representing one mouse. The metabolic inhibitors used were oligomycin (oligo), carbonyl cyanide p-trifluoromethoxyphenylhydrazone (FCCP), 2-deoxyglucose (2-DG) and rotenone with antimycin (rot/a.a). (*C-E*) Isolated colonic crypts were cultured in 25mM or 100mM of glucose for 5 days and levels of metabolites were measured via liquid chromatography-mass spectrometry. Data points represent mean +/− SEM and representative of 1 experiment (n=3). Stats represent student’s t-test with Benjamini-Hochberg procedure, where \**P*<0.05, \*\**P*<0.01, \*\*\**P*<001, \*\*\*\**P*<0001. (*F-L*) Murine colonoids were cultured in 70mM of sucrose, fructose, glucose, or no-sugar-added control, with or without DCA (dichloroacetate). (*F*) Representative images of murine colonoids after 5 days of culture in sugar and metabolic inhibitors. Images taken at 4X magnification (scale bars= 200μm). (*G*) Number of colonoids developed per well with DCA treatment and PBS vehicle control. (*H*) Viability of colonoids cultured with rotenone, DCA and 2-DG is shown. (*I-L*) Colonoids were isolated in Trizol and analyzed via RNAseq. (*I*) PCA plot showing variance across groups with percentages on axes representing percent variance explained by each principle component. (J) Transcript expression level of epithelial subset gene signatures as determined colonoid RNAseq, where red and blue represent high or low expression level, respectively, normalized across rows. *(K*) GSEA of colonoid RNAseq data showing enrichment of genes in Control-treated compared to Glucose treated colonoids and *(L)* Glucose-treated compared to Glucose/DCA-treated for gene sets indicated. Data points represent mean +/− SEM and representative of 2 experiments (n=3) and * represents genes that are significantly differentially expressed in control-versus glucose-treated colonoids (FC>1.5, *P*<0.05, FDR<0.3). Stats represent one-way ANOVA with multiple comparisons to Vehicle, where \**P*<0.05, \*\**P*<0.01, \*\*\**P*<001, \*\*\*\**P*<0001.

### High sugar diet reduces proliferative potential of Lgr5^+^ colonic epithelium

Given the reduction in transcripts associated with Lgr5^+^ ISCs and TA cells, we postulated that HS diet was specifically affecting these critical cells and their proliferative capacity. Using *Lgr5^eGFP-Cre-ERT2^* reporter mice, we isolated Lgr5^+^ stem cells and their immediate daughter cells (which retain low expression of Lgr5) for RNAseq after 2 weeks of HS or HF diet (Nick Barker et al., 2007). Similar to the total epithelium, Lgr5^+^ ISCs from HF-fed mice were only modestly different from ISCs from HS-fed at the level of the whole transcriptome, but showed much more substantial enrichment in the expression of proliferation-related genes targeted by Myc and E2F. In contrast, Lgr5^+^ ISCs from HS-fed mice showed a clear reduction in many of the cell cycle genes that comprise the TA cell gene signature, suggesting that a HS diet reduces the proliferative capacity of ISCs and their daughter cells prior to the induction of damage (Fig. 4 *F* and *G* and Fig. S5 *D* and *E*). Lineage tracing of Lgr5^+^ daughter cells (from *Lgr5^eGFP-Cre-ERT2^/Rosa^LSL-TdTomato^* mice) showed that HS-diet, DSS-treated mice showed reduced cell migration up the crypt wall, as indicated by the distance and relative position of tamoxifen-activated Tomato^+^ cells from GFP^+^ ISCs located at the base of the crypt (Fig. 4 *H* and *I* and Fig S5*F*).

### Excess dietary sugar alters colonic crypt metabolism, increasing spare respiratory capacity and glycolytic response to a glucose challenge

It was unclear how our HS diet might affect the proliferation of ISCs, but altering sugar concentrations can modify cell metabolism and thereby proliferation rates (Palmer, Ostrowski, Balderson, Christian, & Crowe, 2015). To test how a HS diet might affect the metabolism of ISCs and their daughters, we isolated crypts and analyzed their glycolytic and respiratory rates (Fan et al., 2015). Colonic crypts isolated from HS or HF-fed mice only modestly differed in their basal aerobic respiration (oxygen consumption rate; OCR) or anaerobic respiration (extracellular acidification rate; ECAR), but as expected, displayed a high reliance on aerobic respiration as demonstrated by a high OCR:ECAR ratio (Fig. 5*A* and Fig. S6 *A-D*). Crypts from HS-fed mice showed a significant increase in the difference between basal and maximal oxidative rates, termed spare respiratory capacity (SRC; as determined by uncoupling ATP synthesis from the electron transport chain with FCCP to maximize oxygen consumption) (Fig. 5*A*). High SRC levels are associated with high Complex II activity, and likely reflects unused aerobic respiration capacity in crypts from HS-fed mice (Pfleger, He, & Abdellatif, 2015). We next carried out a glucose stress test to measure the glycolytic capacity of crypts isolated from HS or HF-fed mice. This assay begins with 3 hours of glucose deprivation to remove effects of stored glucose. Interestingly, crypts from HS-fed mice were still able to carry out glycolysis even after this period of deprivation, before the addition of exogenous glucose, indicating that a HS diet is leading to a substantial reservoir of sugar within the epithelium (Fig. 5*B*). The increased glycolytic rate of crypts from HS− fed mice was maintained throughout the assay, supporting the idea that dietary sugar directly activated this metabolic pathway in epithelial cells (Fig. 5*B*). After the addition of oligomycin, which blocks the production of ATP from aerobic respiration, neither crypts from HS− nor HF-fed mice exhibited a compensatory increase in glycolysis, as has been measured in most other cells (Fig. 5*B*). In concert with the increased SRC of colonic epithelium from HS-fed mice, the lack of a compensatory increase in glycolysis in colonic crypt cells when respiration is blocked suggests that glycolysis and aerobic metabolism may be uncoupled in colonic crypts and that these cells may not have the capacity to rapidly switch their metabolic profile when nutrient availability changes. In accord, RNAseq analysis of the epithelium revealed a distinctly increased expression of glycolysis-regulating enzymes, such as *Hk2*, *Hk3*, and *Pfkfb3* in HS-fed epithelium treated with DSS (Fig. S6 *E-G*). The intestinal epithelium typically relies on aerobic respiration via fatty acid oxidation (Fan et al., 2015). Thus, we hypothesize that colonic crypts from HS-fed mice are primed for an increased glycolytic rate, yet are unable to efficiently utilize the glycolytic metabolites for respiration, as demonstrated by an increased SRC.

### Coupling glycolysis with aerobic respiration rescues colonoid development when cultured in excess sugar

Given HS diet increased glycolytic potential of crypts without a requisite increase in aerobic respiration, we postulated that HS conditions may be impairing the flux of glucose and its downstream metabolites into the TCA cycle. Utilizing colonoids cultured in high or low glucose conditions, we compared total metabolite pools. In accord with our glucose stress test of isolated crypts, we found that HS-cultured colonoids have greater stores of intracellular-glucose (Fig. 5*C*). HS-cultured colonoids also had higher levels of pyruvate and aspartate, but significantly reduced levels of the TCA cycle metabolite, *α*-ketoglutarate, suggesting reduced conversion of pyruvate to acetyl-CoA and a reduced flux of glucose metabolites into mitochondrial respiration (Fig. 5*D*). Amino acids such as serine, alanine and glycine were increased under HS-conditions, supporting the notion that pyruvate entry into the TCA cycle is inhibited under HS conditions (Fig. 5*E*). RNAseq analysis demonstrated that the epithelium from HS-fed, DSS-treated mice expressed greater levels of the Pyruvate Dehydrogenase kinases (PDHKs) *Pdk1* and *Pdk2*, compared to DSS-treated, HF-fed mice (Fig. S6 *E-G*). When active, Pdk1 and Pdk2 inactivate pyruvate dehydrogenase, blocking the flux of glycolytic metabolites into the TCA cycle by inhibiting the conversion of pyruvate to acetyl-CoA. Therefore, increases in pyruvate PDHKs may explain the inefficient utilization of glycolytic metabolites. To determine whether these PDHKs are responsible for impairing epithelial regeneration, we treated isolated ISCs with dichloroacetate (DCA), a PDHK inhibitor. DCA is a drug used to increase aerobic utilization of glucose by increasing the rate at which pyruvate is converted to acetyl-CoA and enters the TCA cycle (Madhok, Yeluri, Perry, Hughes, & Jayne, 2010; Michelakis et al., 2010; Shahrzad, Lacombe, Adamcic, Minhas, & Coomber, 2010; J. Y. Y. Wong, Huggins, Debidda, Munshi, & De Vivo, 2008). Treatment of developing organoids revealed that DCA significantly increased viability and organoid number developing from ISCs cultured in inhibitory levels (70mM) of sucrose, fructose and glucose (Fig. 5 *F* and *G*). We did not observe similar improvements in organoid development when treating stem cells with rotenone, an inhibitor of the mitochondrial respiratory chain, and only modest improvements with 2-deoxyglucose, a glycolysis inhibitor (Fig. 5*H*). RNAseq analysis of colonoids treated with excess glucose had a significant reduction in the expression of core ISC genes such as *Lgr5*, *Axin2* and *Ascl2*, all of which were substantially restored with DCA treatment (Fig. 5 *I-J*). Further, GSEA showed enrichment in control-treated colonoids for gene sets associated with E2F targets and the Intestinal Stem Cell signature, the latter of which was restored with DCA treatment of glucose-impaired colonoids (Fig. 5 *K* and *L*). Therefore, we hypothesize that it is not high glycolytic rates or reduced mitochondrial activity that is impairing ISC function, but rather a PDHK-mediated deviation of glucose metabolism away from mitochondria and by forcing Lgr5^+^ ISCs and their progeny to utilize glucose aerobically, their proliferative and differentiating capacity was rescued.

## Discussion

We report that excess dietary sugar leads to lethal colonic damage in mice treated with DSS. We demonstrated direct sugar-induced impairment of ISC growth into organized colonoids *in vitro*. Further, after 3 days of DSS treatment, HS− fed mice already exhibited an impaired epithelial proliferative response with reduced Ki67^+^ staining and fewer daughter cells from Lgr5^+^ intestinal stem cells. Failed proliferation of ISCs and TA cells in colonic crypts from HS-fed mice was associated with an increase in glycolytic response to glucose deprivation that did not coincide with a requisite increase in respiration. Accordingly, pyruvate accumulated in colonoids cultured in HS conditions but TCA cycle intermediates were reduced. By biochemically recoupling glycolysis to aerobic respiration, we restored stem cell growth and colonoid development under high sugar conditions, suggesting sugar-induced shifts in metabolism can directly reduce the proliferation of the colonic epithelium.

In contrast to previous studies using diets high in sugar and low in fiber, or supplementing standard chow with sugar water, we did not correlate increased susceptibility to significant shifts in the microbiome (Desai et al., 2016; Khan et al., 2020; Laffin et al., 2019). We cannot rule out the possibility that these taxa may exacerbate disease once inflammation has begun and indeed, the HS-diet-associated microbiome did contribute to worse outcomes in gnotobiotic mice, but altogether our data indicated that intestinal bacteria are unlikely the cause of the massive and rapid failure of the colonic epithelium seen in DSS-treated HS-fed mice. We also did not observe a distinct outgrowth in mucophilic bacteria (*Akkermansia spp*.) in our HS-fed mice (Desai et al., 2016; Khan et al., 2020). We suspect that this is because our HS diet was low in fiber, but not fiber free and thus we may have been able to avoid the expansion of mucus-degrading bacteria.

Sugar is typically absorbed by the duodenum and it is unlikely that our ancestors’ diet contained foods with high enough sugar concentrations to reach the colon (Röder et al., 2014; Spreadbury, 2012; Yoshikawa et al., 2011). However, soft drinks and other modern, processed foods contain large concentrations of acellular sugar and it is possible that sugar reaches the colon when these foods are consumed (Khan et al., 2020). Unlike the small intestine, the colonic epithelium does not express the bi-directional glucose transporter GLUT2 (Wang et al., 2020), that transports glucose into the bloodstream and we hypothesize that the colonic epithelium is a terminal endpoint for sugar, making it more susceptible to increases in dietary sugar. Indeed, even after 3 hours of glucose deprivation, HS-fed crypts were still able to perform glycolysis demonstrating glucose storage capacity in these cells. One prediction from this hypothesis is that regeneration of the small intestine would be much less affected by high sugar concentrations because the small intestinal cells pass sugar to the host.

We found that high sugar concentrations prevented the development of colon-derived organoids from stem cells but was not toxic to established colonoids. This indicates that sugar is likely directly impairing ISCs function or the earliest progenitor cells, rather than impacting differentiated epithelium. ISCs have distinct metabolic needs compared to most other cell types. For example, in the small intestine, Paneth cells have been shown to metabolically tune ISCs, performing glycolysis to produce lactate before transferring it to ISCs, where it is converted to pyruvate and used for respiration (Rodríguez-Colman et al., 2017; Sato et al., 2011). That ISCs require an adjacent cell to carry-out glycolysis supports the hypothesis that this metabolic pathway may have negative effects on their biology. Further, ISCs with impaired fatty acid oxidation leads to disrupted self-renewal and ultimately loss of Lgr5+ stem cells (Chen et al., 2020) demonstrating the importance of fuel metabolism in ISC function.

Based upon our finding of increased glycolytic response to glucose deprivation and increased SRC in colonic crypts of HS-fed mice, we postulate that ‘forced’ glycolysis may actually impede aerobic respiration in ISCs and their adjacent daughter TA cells. One potential mechanism for this phenotype is regulation of glycolysis and cell cycle via pyruvate dehydrogenase kinases (PDHKs) in Lgr5^+^ cells, which inhibits pyruvate conversion to acetyl-CoA and further catabolism via the TCA cycle. Interestingly, in hematopoietic stem cells, PDHKs are important in reducing glycolytic activity and promoting quiescence (Takubo et al., 2013). We hypothesize that PDHKs have much the same function in intestinal stem cells and, when they are increased by high sugar concentrations, they prevent regenerative proliferation in the crypt.

Intestinal regeneration is necessary to maintain barrier integrity, especially in patients exposed to direct intestinal damage such as flares of IBD or radiation therapy. Here we have shown the deleterious effects of consuming excess dietary sucrose in a murine model of intestinal damage. Treatment of active flares of Crohn’s Disease and Ulcerative Colitis often involve exclusive enteral nutrition which can contain high amounts of sugar and emulsifiers (Grover, Muir, & Lewindon, 2014). Numerous studies, including our own, have now shown the negative impact of high sugar diet in murine models of colitis, suggesting that we may improve these therapies by reducing sugar content (11–13). Indeed, both murine studies and clinical trials in pediatric cohorts have found that diets lower in sugar lead to better outcomes in patients that exhibited intestinal inflammation (Obih et al., 2016; Yeh et al., 2019). Therefore, in order to better treat patients exhibiting high levels of intestinal damage, whether it be from infection, auto-inflammation, or radiation, it is imperative that we better understand how different dietary components may impact the regenerative capacity of the intestinal epithelium.

## Materials and Methods

### Mouse Models and Treatments

5-week-old wild type *C57BL/6Tac* mice (B6 MPF; Taconic) were used for diet and DSS treatment unless otherwise noted. *Lgr5^eGFP-Cre-ERT2^* (008875) were bred with *Rosa^TdTomato^* (007909) both purchased at Jackson laboratories. Both male and female age-matched mice (5-8 weeks) were used for all experiments. All experiments were performed in an American Association for the Accreditation of Laboratory Animal Care-accredited animal facility at the University of Pittsburgh. Mice were kept in specific pathogen-free conditions and housed in accordance with the procedures outlined in the Guide for the Care and Use of Laboratory Animals under an animal study proposal approved by the Institutional Animal Care and Use Committee of the University of Pittsburgh. The number of mice used in each experiment was determined given a predicted effect size of 30% in experimentally measured variables (number of cells, weights and survival) in DSS-induced colitis would show significance and the variance within groups could be 15%. In order to have a power of 0.8 and a probability error of 0.05 or less, we used 4 mice per group and 3 repeats for a total of n=12 to reach sample size large enough to detect differences. We expected a 2-fold difference of expression and metabolite accumulation in our enteroid model, given our preliminary data. Therefore, we used tissue from three independent samples for each experiment type to achieve a power of 0.8 and a probability error of 0.05 or less. Upon arrival from Taconic, mice were placed on two special diets (Envigo, Madison, WI) with kilocalories consisting of 18% protein (casein and methionine), 12% fat (soybean oil), and 70% carbohydrates. The High Sugar (HS, TD.160477) diet derives 94% of carbohydrates from sucrose and High Fiber diet (HF, TD.160476) contains 94% of carbohydrates from high amylose cornstarch. Standard diet was composed of 26.1% protein, 59.6% carbohydrates and 14.3% fat by Kcal (Prolab IsoPro RMH 3000, 5P75). Mice were provided food *ad libitum* for 2 weeks and then provided dextran sodium sulfate (DSS) at 3% by weight in their drinking water *ad libitum* for 1 week. Weights were taken twice weekly during the initial diet change phase and daily once DSS was initiated and for one week after changing DSS water back to untreated water. Gnotobiotic C57BL/6 female 8-week mice were housed in germ-free conditions and then gavaged with the microbiome of HS or HF-fed mice, but kept on standard facility chow in separate isolators. After 3 days of colonization, mice were started on 3% DSS *ad libitum* and weights were taken daily. Water supplemented with SCFA contained sodium acetate (0.554g/100mL), sodium butyrate (0.441g/mL) and sodium propionate (0.249g/mL). Tributyrin was added to high sugar food (5% by weight) and glycerol was added to high sugar and high fiber food (5% by weight) as controls.

### Histological analysis of colonic tissue

Distal colon samples were fixed in formalin, dehydrated and paraffin embedded. Sections were stained with hematoxylin and eosin (H&E) stains for morphological analysis and by TUNEL staining for apoptotic cell detection. Histopathology analysis was blinded and determined following the scoring criteria: 1) degree of inflammation in lamina propria (score 0-3); 2) loss of goblet cells (score 0-2); 3) abnormal crypts or epithelial hyperplasia with nuclear changes (score 0-3); 4) presence of crypt abscesses (score 0-3); 5) mucosal erosion and ulceration (score 0-1); 6) submucosal spread to transmural involvement (score 0-3) and 7) number of neutrophils (score 0-4). Scores for the seven parameters were combined for a total maximum score of 17 (Ostanin et al., 2009). Distal colon samples were also fixed in Carnoy’s fixation, dehydrated and paraffin embedded and sections were stained with Alcian blue and PAS stain for mucus detection. Quantification of TUNEL was measured using ImageJ software analysis.

### 16s rRNA gene analysis of bacterial abundance in intestine

Fecal samples were collected on the first day mice were started on their diets (Initial), 2 weeks after starting their diets (Standard or Defined Diets) and during DSS treatment (DSS). DNA was isolated using the MoBio Power Soil Isolation Kit and PCR amplified at the V4 region of the 16S rRNA gene (515F-806R) and sequenced at the Argonne National Library on an Illumina MiSeq instrument. Microbiome informatics were performed using QIIME2 2020.2 (Bolyen et al., 2019). Raw sequences were quality-filtered and denoised with DADA2 (Callahan et al., 2016).

Amplicon variant sequences (ASVs) were aligned with mafft and used to construct a phylogeny with fasttree2 (Katoh, Misawa, Kuma, & Miyata, 2002; Price, Dehal, & Arkin, 2010). Alpha diversity metrics (observed OTUs), beta diversity metrics (Bray Curtis dissimilarity) and Principle Coordinate Analysis (PCoA) were estimated after samples were rarefied to 63,000 (subsampled without replacement) sequences per samples. Taxonomy was assigned to ASVs using naive Bayes taxonomy classifier against the Greengenes 18_8 99% OTUs reference sequences (McDonald et al., 2012). All plots were made with publicly available R packages.

### Blood glucose assay

Mice were fed defined diets for 2 weeks then either fasted or allowed to eat overnight and blood was taken from the retro-orbital sinus after anesthesia with isofluorane. Glucose levels were measured using a Precision Xtra glucometer.

### FITC-dextran assay

To evaluate gut permeability, 4kDA FITC-dextran (Sigma-Aldrich) was dissolved in PBS (100mg/ml) and mice were orally gavaged at 44mg/100g of body weight after fasting for 8 hours. Mice were euthanized and blood was collected immediately via cardiac puncture. Serum was isolated and diluted with an equal volume of PBS, of which 100μL was added to a 96-well microplate in duplicate. The plate was read at an excitation of 485nm and an emission wavelength of 528nm to quantify FITC in blood, using a serially dilutes FITC-dextran to calculation concentration. Mice treated with DSS on standard diet and gavaged with FITC-dextran were used as a positive control of mice with a damaged intestinal barrier.

### Gene expression profiling by RNAseq and bioinformatics analyses

Bulk epithelium was isolated from Rag1−/− mice by scraping the apical side of the colonic tissue to release cells and placing in trizol to isolate RNA. Mice were fed defined diets for 2 weeks and were either untreated (n=3) or treated with 3 days of 3% DSS drinking water (n=4). DSS treated samples were precipitated overnight in Lithium Chloride to remove DSS that may interfere with the sequencing process. Lgr5^+^ cells were isolated from the colons of Lgr5^eGFP-IRES-Cre-ERT2^ reporter mice fed defined diets for 2 weeks, as described previously with some modifications (Fan et al., 2015). Briefly, colons were butterflied and vortexed to remove luminal contents then incubated at 37°C for 30 minutes in EDTA to dissociate the epithelium from the lamina propria. Vortexing released crypts, which were passed through a 20-gauge needle to dissociate further into single cell suspension as well as passed over a 20-micron filter to further break up any remaining clumps of cells. Cells were stained with a Live/Dead discrimination dye and antibodies against EPCAM and CD45.2 and then resuspended in rock-inhibitor containing DMEM to prevent differentiation of Lgr5+ stem cells. Live cells were sorted on the MoFlo Astrios (Beckman) cell sorter directly into Takara kit lysis buffer (SmartSeq HT). Cultured colonoids were grown in the conditions listed below and their RNA was extracted via Trizol separation. DNA libraries were prepared (Nextera XT kit) and RNA-sequencing was performed on Illumina NextSeq500 by the University of Pittsburgh Health Sciences Sequencing Core. Adapter sequences were trimmed from raw reads using Trimmomatic with default parameters.

TopHat2.1.1 was used to map trimmed reads onto mouse genome build mm10 and Cufflinks was used to calculate gene expression values (FPKM; fragments per kilobase exon per million mapped reads) (Bolger, Lohse, & Usadel, 2014; Trapnell et al., 2012). Enrichment of genesets were calculated using Gene set enrichment analysis (GSEA) from the Broad Institute (http://www.broad.mit.edu/gsea). Heatmaps were created using Morpheus from the Broad Institute (https://software.broadinstitute.org/morpheus) from FPKM log2 transformed expression levels.

### Tamoxifen (TX) administration

Tamoxifen (TX, Sigma-Aldrich), was orally gavaged at 5mg/mouse/day, on the first day of DSS treatment. Since TX is poorly soluble in water, the amount needed for a single day was dissolved in 95% ethoanol with heating to 37°C and then diluted in corn oil (Sigma) such that 100μL had 5mg.

### Microscopy

Distal colonic tissue was flushed of luminal contents using PBS and then fixed for 1 hour in 2% PFA, dehydrated in 30% sucrose overnight and flash-frozen in OTC media. Sections were stained with antibodies specific to EPCAM (BioLegend, clone G8.8, catalog # 118212) and Ki67 (invitrogen, clone SolA15, ref 14-5698-82) overnight and 5 minutes for the Hoechst nuclear stain for (Invitrogen, ref H3570). Images were taken on Zeiss LSM 510 and Nikon A1 confocal microscopes and analyzed using ImageJ software.

### Flow cytometry

All antibodies used for flow cytometry were purchased from either ThermoFisher, BD Biosciences, or BioLegend. The antibodies we used for flow cytometry are: CD45.2 (Invitrogen, clone 104, ref 47-0454-82), EPCAM (BD Biosciences, clone G8.8, catalog # 563478), activated-Caspase-3 (BD Biosciences, clone C92-605, catalog # 560901), and Ki67 (BioLegend, clone 16A8, catalog # 652403). Dead cells were discriminated in all experiments using LIVE/DEAD fixable dead stain (ThermoFisher, catalog # 501121526). All stains were carried out in media containing anti-CD16/32 blocking antibody (ThermoFisher, clone 93, catalog # 14-0161-86). All flow cytometry was acquired on an LSRFortessa FACS analyzer. Cells were isolated from the colon for flow cytometry using EDTA and DTT dissociation and shaking to release the epithelium from the lamina propria (Hall et al., 2011). To separate intraepithelial cells, the cell suspension was spun down in a 30% percoll gradient. Analysis of flow cytometry was carried out on FlowJo software (TreeStar).

### Seahorse Metabolic Flux Analysis

Crypts were isolated as described previously (25). Crypts were seeded at 150crypts/50μL on Cell-Tak-coated Seahorse Bioanalyzer XFe96 culture plates (300,000 or 100,000 cells/well, respectively) in assay media consisting of minimal, unbuffered DMEM supplemented with 1% BSA and 25 mM glucose, 2 mM glutamine, and for some experiments, 1 mM sodium pyruvate and Matrigel. Basal rates were taken for 30 min, and in some experiments, oligomycin (2 μM), carbonyl cyanide p-trifluoromethoxyphenylhydrazone (FCCP) (0.5 μM), 2-deoxy-d-glucose (10 mM), and rotenone/antimycin A (0.5 μM) were injected to obtain maximal respiratory and control values. Spare respiratory capacity (SRC) was measured as the difference between the basal oxygen consumption rate (OCR) and the maximum OCR after FCCP injection. A glucose stress test was used to determine glycolytic response of crypts, where crypts were placed in glucose free media for 3 hours prior to adding exogenous glucose and extra cellular acidification rate (ECAR) values were measured while oligomycin (2 μM), 2-deoxy-d-glucose (10 mM), and rotenone/antimycin A (0.5 μM) were injected to wells. Figure panels show a representative trace of one experiment and combined data for SRC and glycolytic rate (calculated as the difference between maximal ECAR after glucose injection and basal ECAR after 2-DG injection).

### Fluorescent glucose tracing

Mice were fasted for 8 hours and then anesthetized using isofluorane and a murine colonoscope (Storz) was inserted. To remove luminal fecal contents and mucus, the colon was rinsed using 200μL of PBS. After a period of 30 minutes, mice were woken up to allow any fecal matter to evacuate and then anesthetized again to introduce 100μL of Cy-5-labelled glucose or Cy-5 secondary goat anti-rat antibody (0.1mM diluted in PBS, ThermoFisher A10525) into the colon via a gavage needle enema. Mice were supported inverted for 1 minute after the enema to ensure the probe remained in the colon and then were taken down 30 minutes after and distal colon samples were collected, fixed in 2% PFA and dehydrated in 30% sucrose overnight.

### Mouse crypt-derived organoid generation

Mouse intestinal crypt-derived organoids were generated as described previously (Beyaz et al., 2016). Briefly, 8-12 weeks old mice were euthanized in a CO2 chamber. The whole intestine was extracted and cleaned from fat, connective tissue, blood vessels and flushed with ice-cold PBS. The intestine was cut into smaller pieces after lateralization and incubated in 7.5 mM EDTA in ice-cold PBS with mild agitation for 45 minutes at 4 C. Then, the crypts were mechanically dissociated from tissue via shaking and strained through a 40-micron strainer.

After washing with ice-cold PBS and centrifugation at 300 r.c.f. for 5 minutes in a microcentrifuge (Thermo Fisher 0540390), isolated crypts were counted and embedded in Matrigel (Corning 356231 Growth Factor Reduced) in 1:4 ratio at 5-10 crypts per μL and plated in 24-well plates (25 ul dome/ well). The Matrigel was allowed to solidify for 8-15 minutes in a 37C incubator and solidified domes were cultured in Advanced DMEM (Gibco) media supplemented with recombinant murine Chiron 10 μM (Stemgent), Noggin 200 ng ml−1 (Peprotech), R-spondin 500 ng ml−1 (R&D or Sino Biological), N2 1X (Life Technologies), B27 1X (Life Technologies), Y-27632 dihydrochloride monohydrate 20 ng ml−1 (Sigma-Aldrich), EGF 40 ng ml−1 (R&D), N-acetyl-L-cysteine 1μM (Sigma-Aldrich). 500μL of crypt media was changed every other day and maintained at 37C in fully humidified chamber containing 5% CO2.

### Mouse organoid propagation

Organoids were propagated by dissociating crypt-derived organoids in TryplE Express (Invitrogen) for 3min at 37C. After this time, the TryplE Express was quenched by adding 1-2x that amount of Advanced DMEM/F12 (Gibco). The pellet containing the dissociated intestinal single cells after centrifugation in a microcentrifuge (Thermo Fisher 0540390) at 300 r.c.f. for 5min was resuspended in Matrigel (Corning 356231 Growth Factor Reduced) and embedded onto a flat bottom 24 well cell culture plate (Corning 3526) by forming 20μL droplets of Matrigel, creating at least three technical replicates for each condition. The embedded Matrigel droplets were immediately placed inside a fully humidified incubator containing 5% CO2, which was maintained at 37C for 5min to solidify the Matrigel droplets. Once the Matrigel was solidified, 600μL of supplemented Advanced DMEM/F12 cell medium described above was added to each well. The media was changed every 2 days for each well and the plate was maintained in a 37C incubator.

### Human patient-derived colon organoid generation

Human colon organoids were generated as described previously with minor modifications (Beyaz et al., 2016). Briefly, normal colon tissue samples were obtained from patients with informed consent undergoing surgical resection procedures at Northwell Health. Study protocols were reviewed and approved by the Northwell Health Biospecimen Repository (NHBR-1810). Tissue samples were first cut into small pieces, about 0.5cm² and incubated at 4C in an antibiotic mixture consisting of 1X PBS +100ug/mL Normocin (Invivogen Cat# ant-nr-1), 50ug/mL Gentamicin (Amresco, Cat# E737), and 1X Pen/Strep (ThermoFisher Cat# 15070063) for 15 minutes. Next, the pieces were washed with 1X PBS before a 75-minute incubation in a 5mM EDTA solution at 4C on a rocker. After incubating, the tissue samples were washed once more with 1X PBS. Crypts were then released from the tissue by shaking the pieces in a tube with ice cold 1X PBS. Crypts in the supernatant were transferred to a new tube and spun down at 100g for 5 minutes at 4C. These isolated crypts were then embedded in a 70/30 Matrigel (Corning, Cat# 356231) and culture medium mixture and plated in 40μL droplets on 12 well plates. The Matrigel was allowed to polymerize at 37C for 15 minutes before adding 1mL of culture medium to each well, with the culture medium consisting of Advanced DMEM (Life Technologies 12634028), 1X Glutamax (Life Technologies 35050061), 10mM HEPES (Thermo Fisher Scientific 15630080), 50% WRN conditioned medium (Homemade), 1X B27 (Life Technologies 12587010), 1X N2 (Life Technologies 17502048), 10mM Nicotinamide (Sigma Aldrich N0636), 1mM N-acetyl cysteine (Sigma Aldrich A9165), 100 μg ml−1 Primocin (Invivogen ant-pm-1), 10μM SB202190 (Sigma Aldrich S7067), 10μM Y-27632 (Tocris 1254), 10nM Gastrin I (Sigma Aldrich G9020), 50 ng ml−1 EGF (Peprotech AF-100-15), and 500nM A83-01 (Sigma Aldrich SML0788).

### Human patient-derived colon organoid propagation

Human colon organoids were dissociated in Cell Recovery Solution (Corning 354253 Growth Factor Reduced) for up to one hour at 4C. Once the Matrigel was dissolved, the organoids were spun at 500 r.c.f. for 5 minutes at 4C. The supernatant was removed and the pellet was resuspended in TryplE Express (ThermoFisher 12604039). Following a 5 minute incubation at 37C, the digestion was stopped by adding Advanced DMEM. The solution then was centrifuged at 500 r.c.f. for 5 minutes at 4C. Dissociated cells were seeded in 40μL Matrigel droplets and culture medium mixture. Culture medium was then added to each well after the domes polymerized.

### Sugar dose response in organoids

Dose response experiments using organoids were carried out using D-(-)-Fructose (Sigma-Aldrich F0127), D-(+)-Glucose (Sigma-Aldrich G7528), Sucrose (Sigma-Aldrich S0389) at 200, 100, 50, 25, 12.5, 3.1 and 0.8 mM concentrations. Briefly, normal colon organoids were dissociated to near single cells and plated onto a 24 well plate, with 20μL domes per well. 500μL of culture medium further supplemented with different concentrations of D-(-)-Fructose (Sigma-Aldrich F0127), D-(+)-Glucose (Sigma-Aldrich G7528), or Sucrose (Sigma-Aldrich S0389) was added to each well at time of seeding. Media with the supplemented sugars was refreshed every 2-4 days and growth was followed up to day 12. Alternatively, normal colon organoids were dissociated to near single cells and plated onto a 24 well plate, with 20μL domes per well. 500μL of standard culture medium was added and organoids were allowed to grow to D5. Media was then changed to media with supplemented glucose, fructose, or sucrose at varying concentrations. Growth was followed for the next 48 hours.

### Murine intestinal organoid intracellular metabolite isolation

Unless otherwise stated, the dissociation, centrifugation, embedding of Matrigel droplets, media addition, and incubator use are the same as indicated previously above for the murine intestinal organoid culture. For metabolite analysis, organoids were dissociated as described above and cells were seeded in a 1.5mL 10% Matrigel/90% mouse organoid culture medium slurry for each condition in separate 24 well plates. The medium for the low glucose condition was supplemented with 25mM Glucose (Sigma G7021) and the medium for high glucose condition was supplemented with 100mM Glucose. These plates were spun at 100G for 1min at 4C in a centrifuge (Eppendorf 022623508) to allow settling of cells to the bottom of the wells. Then, both plates were placed in a fully humidified 37C incubator.

On day 3 of culture, the medium in each well was discarded and replaced with 25mM or 100mM C-13 isotopic glucose (Cambridge Isotope Labs CLM-1396) supplemented murine organoid medium. For negative control, empty wells were filled with 1.5mL of the 10% Matrigel/90% murine organoid medium supplemented with 25mM or 100mM C-13 isotopic glucose. 24 hours after tracer incubation, organoid medium from each well per condition and the blank wells were placed into Eppendorf tubes and snap-frozen in liquid Nitrogen. 800μL of 1X PBS was used to mix and collect the organoids from each experimental well into Eppendorf tubes and these were spun at 300G for 1min at 4C. After centrifugation, each tube containing the organoid pellet had its supernatant aspirated without disturbing the pellet. 1mL of metabolite extraction solution (stored at −80C) consisting of 50% Methanol stock (Sigma 322415), 30% Acetonitrile stock (Fisher A998N1), 20% distilled water was added to each pellet and mixed thoroughly. Once mixed, each tube containing the extraction solution was snap frozen in liquid Nitrogen, thawed, and then snap frozen in liquid Nitrogen again.

The thawed tubes were mixed at maximum speed on a thermomixer (Eppendorf 5382000023) set to 4C for 15min. After mixing, they were incubated overnight at −80C. The next day, these tubes were centrifuged for 10min at maximum speed at 4C and the supernatant was collected into Eppendorf tubes while the pellets were kept on ice. This collected supernatant was then centrifuged for 10min at maximum speed at 4C and its supernatant was decanted into autosampler vials (Sigma 29659-U) that were then incubated at −80C until metabolite analysis. 300μL of 0.1M NaOH was added to each tube containing an organoid pellet and was mixed thoroughly followed by max speed incubation for 15min at 4C on a Thermomixer (Eppendorf). These tubes were then centrifuged at 300G for 5min at 4C and the supernatants were used for protein quantification using the DC protein quantification assay protocol (Bio-rad 5000112). After incubation at −80 °C for 1 hour, samples were centrifuged to remove the precipitated proteins and insoluble debris. The supernatants were collected and stored in autosampler vials at −80°C until analysis.

Samples were randomized to avoid bias due to machine drift, and processed blindly. LC-MS analysis was performed using a Vanquish Horizon UHPLC system coupled to a Q Exactive HF mass spectrometer (both Thermo Fisher Scientific). Sample extracts were analyzed as previously described (Mackay, Zheng, van den Broek, & Gottlieb, 2015). The acquired spectra were analyzed using XCalibur Qual Browser and XCalibur Quan Browser software (Thermo Fisher Scientific) by referencing to an internal library of compounds.

### Sugar and inhibitor treatment in murine intestinal organoids

For each chemical’s dose response, the following chemicals were added into the medium for each well depending on the dose used for that well for a total of 600μL cell medium/chemical mix around each dome: Rotenone (Sigma R8875), 2-Deoxy-D-Glucose (Sigma D8375), and Sodium Dichloroacetate (Sigma 347795). Formed organoid numbers were quantified and doses were chosen on the third day in culture.

Unless otherwise stated, the dissociation, centrifugation, embedding of Matrigel droplets, media addition, and incubator use are the same as indicated previously above for the sugar and inhibitor culture. Glucose (Sigma G7021), Sucrose (Sigma S9378), and Fructose (Sigma F0127) as well as the previously stated drugs were added into the cell medium (250μL for each well) of a flat bottom 48 well culture plate (Corning 3526) containing secondary intestinal cells. The chosen concentrations for each chemical are as follows: 150mM Glucose, 150mM Fructose, 150mM Sucrose, 7.8nM Rotenone, 1mM 2-Deoxy-D-Glucose, and 4mM Sodium Dichloroacetate. After embedding the Matrigel domes and solidification, cell medium containing these doses of chemicals was added with three technical replicates per condition including a control containing only the supplemented Advanced DMEM/F12 stated previously. From time zero to time 6 hours, the wells that are supposed to have sugars and inhibitors added together had only the inhibitors added to inhibit the cells and the wells that are supposed to have only sugars added had the control supplemented media added. After 6 hours, the sugars were added alone and with inhibitors to those respective wells. The media for this plate was also changed every two days. After six days in culture, organoid numbers and sizes were quantified and CellTiter-Glo® (CTG) values were obtained and plotted as a percent of the control luminescence (Luminescent Cell Viability Assay, G7570).

### Statistical analysis

Statistical tests used are indicated in the figure legends. Lines in scatter plots represent mean for that group. Group sizes were determined based on the results of preliminary experiments. Mouse studies were performed in a non-blinded manner. Statistical significance was determined with the two-tailed unpaired student’s t-test when comparing two groups or one-way ANOVA with multiple comparisons, when comparing multiple groups, except in the event that there were missing values due to death (e.g. weight loss curves) in which case multiple t-tests were used to compare against the High Sugar diet, Standard diet, or Control group, as stated in the figure legends. All statistical analyses were calculates using Prism software (GraphPad). Differences were considered to be statistically significant when *P*<0.05.

## Data Availability

All relevant data, associated protocols and materials are present in SI.

## Author Contributions

Conceptualization, A.H.P.B., J.J., G.M.D., S.B. and T.W.H.; Formal Analysis, A.H.P.B, A.S.H.C., A.V.M., N.R., C.W.M., A.O.D., A.C.P. and V.S.C.; Investigation A.H.P.B., P.C. J.J., L.M., A.V.M., B.Y., O.E. and K.O.; Resources, S.B., G.M.D and T.W.H.; Original Draft, A.H.P.B. and T.W.H; Writing-Review & Editing, A.H.P.B., S.B., G.M.D. and T.W.H.; Supervision, T.W.H.

## Acknowledgments

The authors would like to thank M. Meyers from UPMC Hillman Institute of Cancer flow core for cell sorting, J. Toothaker for assistance in image analysis, Ece Kilic for assistance with the metabolomics analysis, the staff of the Division of Laboratory Animal Services for the animal husbandry, the UPMC Children’s Hospital of Pittsburgh Histology Core, W. MacDonald and R. Elbakri at the Univ. of Pittsburgh Health Science Sequencing Core, the Center for Biological Imaging, the University of Pittsburgh Center for Research Computing, and the members of the Hand, Beyaz, Delgoffe, Cooper and Poholek labs for helpful discussions. This work was supported by the RK Mellon Institute for Pediatric Research, the NIH (T32AI089443-10 to A.H.P.B.), the Damon Runyon Cancer Research Foundation Postdoctoral Fellowship (2360-19 to A.O.D.) and the Kenneth Rainin Foundation (Innovator’s Award). This work was performed with assistance from the CSHL Mass Spectrometry Shared Resource, which is supported by the Cancer Center Support Grant (5P30CA045508).The authors declare no competing interests.

## Supplemental Figures and Legends

**Table S1.** Dietary composition of standard and defined diets. High fiber and high sugar diets were designed to have the same macronutrient composition (percent calories coming from protein, carbohydrates and fat are kept constant) with identical ingredients. Units indicate gram of ingredient per kilogram of food. * Indicates different ingredients were used in Standard diet (Prolab IsoPro RMH 3000, 5P75).

| Ingredient | Standard | High Fiber | High Sugar |
| --- | --- | --- | --- |
| **Protein** | 26.1% kcal | 17.8% kcal | 17.8% kcal |
| Casein | * | 155.5 g/Kg | 200.0 g/Kg |
| Methionine | 5.8g/Kg | 3.0 g/Kg | 3.0 g/Kg |
| **Carbohydrate** | 59.6% kcal | 70.5% kcal | 70.5% kcal |
| Sucrose | 14.1g/Kg | 22.0 g/Kg | 663.5 g/Kg |
| High-amylose corn starch | 313g/Kg | 699.0 g/Kg | 2.0 g/Kg |
| Maltodextrin | 0g/Kg | 20.0 g/Kg | 20.0 g/Kg |
| Cellulose | 208g/Kg | 9.89 g/Kg | 9.89 g/Kg |
| **Fat** | 14.3% | 11.7% kcal | 11.7% kcal |
| Soybean | * | 39.0 g/Kg | 50.0 g/Kg |
| Vitamin Mix | * | 10.0 g/Kg | 10.0 g/Kg |
| Choline Bitartrate | * | 2.5 g/Kg | 2.5 g/Kg |
| TBHQ, antioxidant | * | 0.01 g/Kg | 0.01 g/Kg |
| Mineral Mix | * | 35.0 g/Kg | 35.0 g/Kg |
| Calcium phosphate, dibasic | * | 4.0 g/Kg | 4.0 g/Kg |

**Fig. S1.**
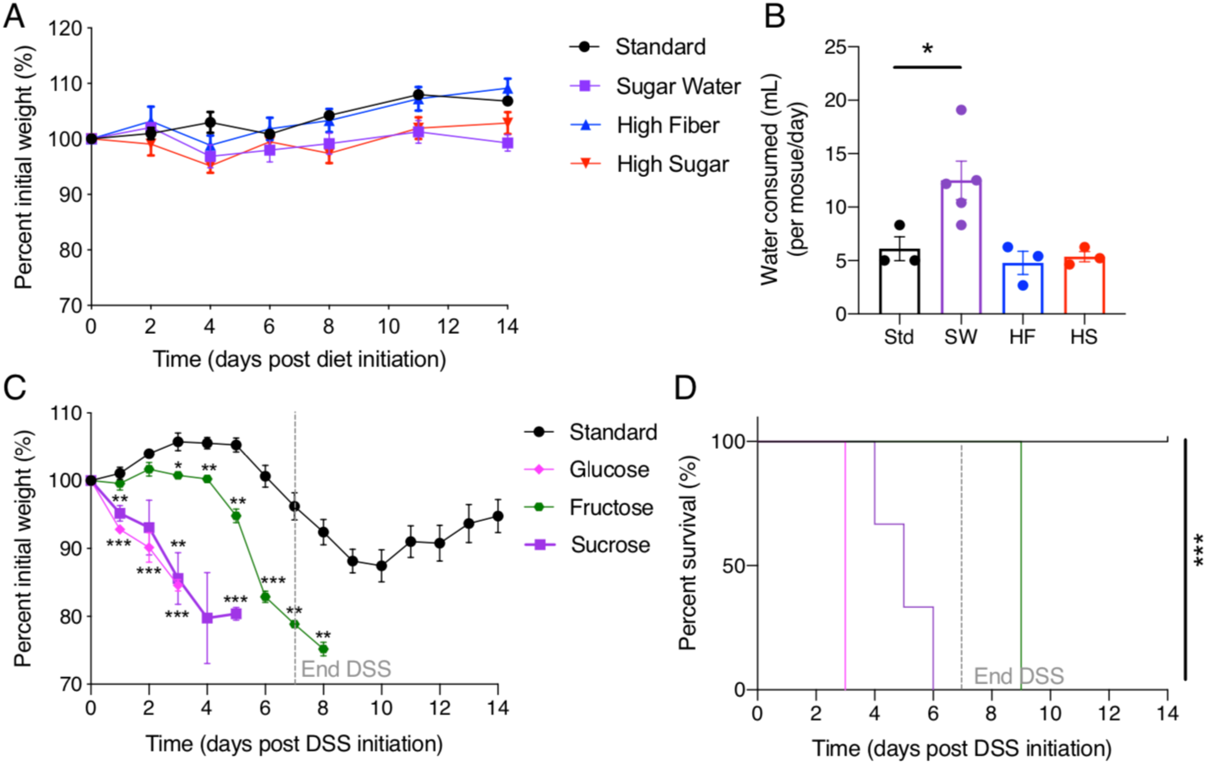
Sugar supplemented water leads to worse DSS-colitis. (*A-B*) Mice were fed high sugar (HS), high fiber (HF), standard (Std) or standard diet with 10% sucrose-supplemented water (SW) for 2 weeks. (*A*) Weight change and (*B*) volume of water consumed on respective diets were measured. Data points represent mean +/− SEM and are representative of 2 experiments (n=3-4). One-way ANOVA used to determine significance in (*B*) where \**P*<0.05. (*C-D*) Mice were fed standard diet and water containing 10% sucrose, glucose or fructose and then treated with 3% DSS in drinking water (dotted line) for one week. (*C*) Percent initial weight and (*D*) survival are shown. Data are presentative of one experiment (n=3). Data points represent mean +/− SEM. Multiple t-tests performed against Standard per day where *\*P<*0.05, *\*\*P<*0.01, *\*\*\*P<*0.001.

**Fig. S2.**
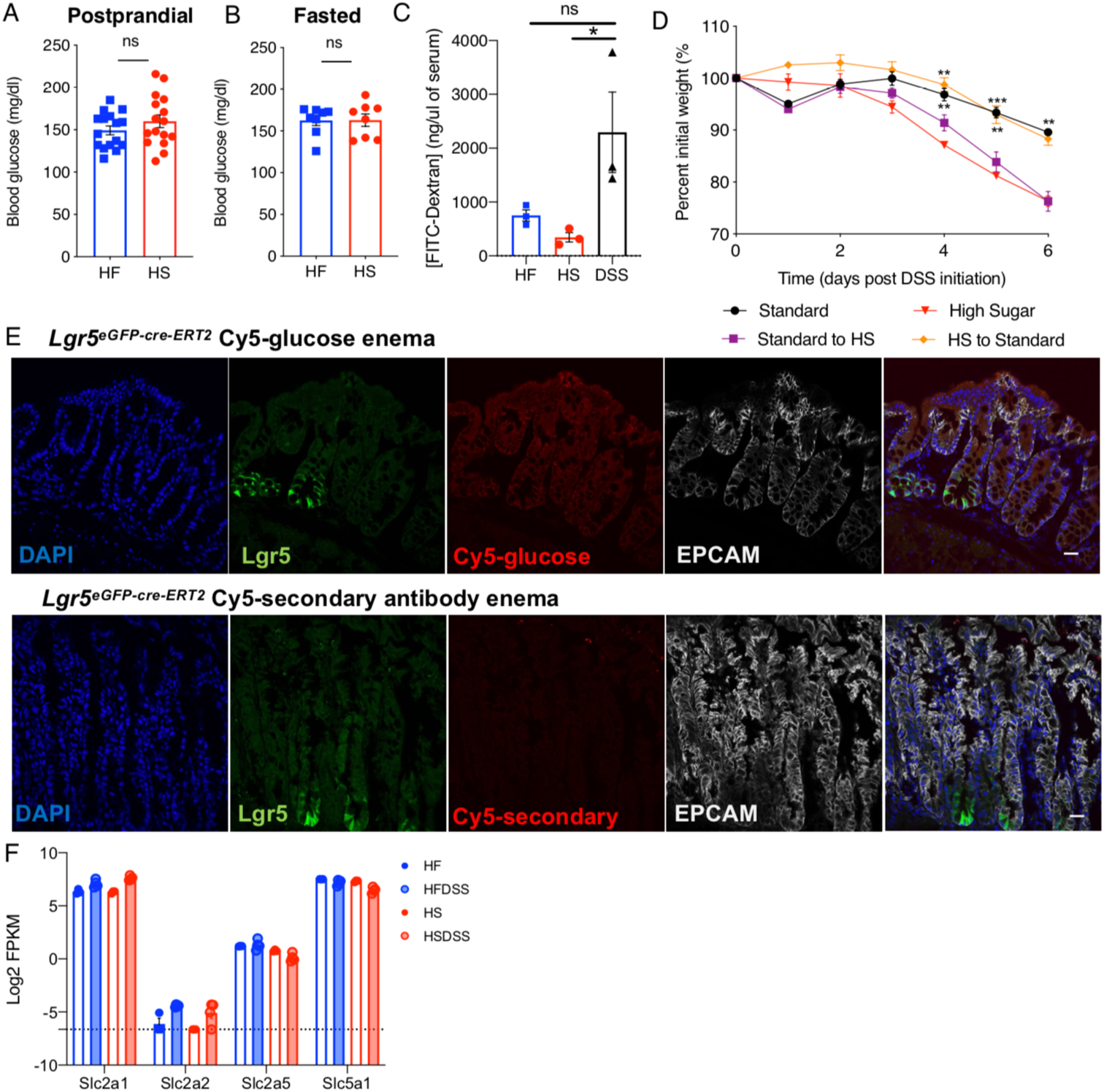
Short-term HS diet does not increase blood sugar or intestinal permeability and luminal sugar must be present during damage. (*A-B*) Blood glucose concentrations of HS or HF-fed mice, (*A*) post-prandial (*B*) and fasted. Data are representative of two to three independent experiments (n=3-4). Each data point represents individual mouse, error bars represents SEM. (C) FITC-dextran recovered from serum of HS or HF-fed mice. Data are representative of two independent experiments (n=3) data points represent individual mouse, error bars represent SEM. One-way ANOVA used to determine significance where *\*P<*0.05. (D) Mice were fed 2 weeks of standard diet and switched to HS diet on the first day of DSS treatment (purple) or fed 2 weeks of HS diet then switched to standard on the first day of DSS treatment (orange). Percent initial weight is shown. Data are representative of 2 experiments (n=3-4) and data points represent mean +/− SEM. Multiple t-tests performed against HS per day where *\*P<*0.05, *\*\*P<*0.01, *\*\*\*P<*0.001. (E) Representative images of colonic sections from fasted Lgr5^IRES-GFP-cre-ERT2^ given a Cy5-glucose enema for 30 minutes prior to sacrifice. Individual and merged channels are shown. Image below is from colonic section of mouse given anti-rat Cy-5 secondary enema as a staining control. Images were taken at 40X magnification, scale bar represents 20μm. (F) Glucose transporter expression level in HS or HF-fed mice with or without 3 days of 3% DSS treatment from RNAseq of colonic epithelium (n=3-4). Data represent individual mouse and error bars represent SEM. Dotted line represents no transcript.

**Fig. S3.**
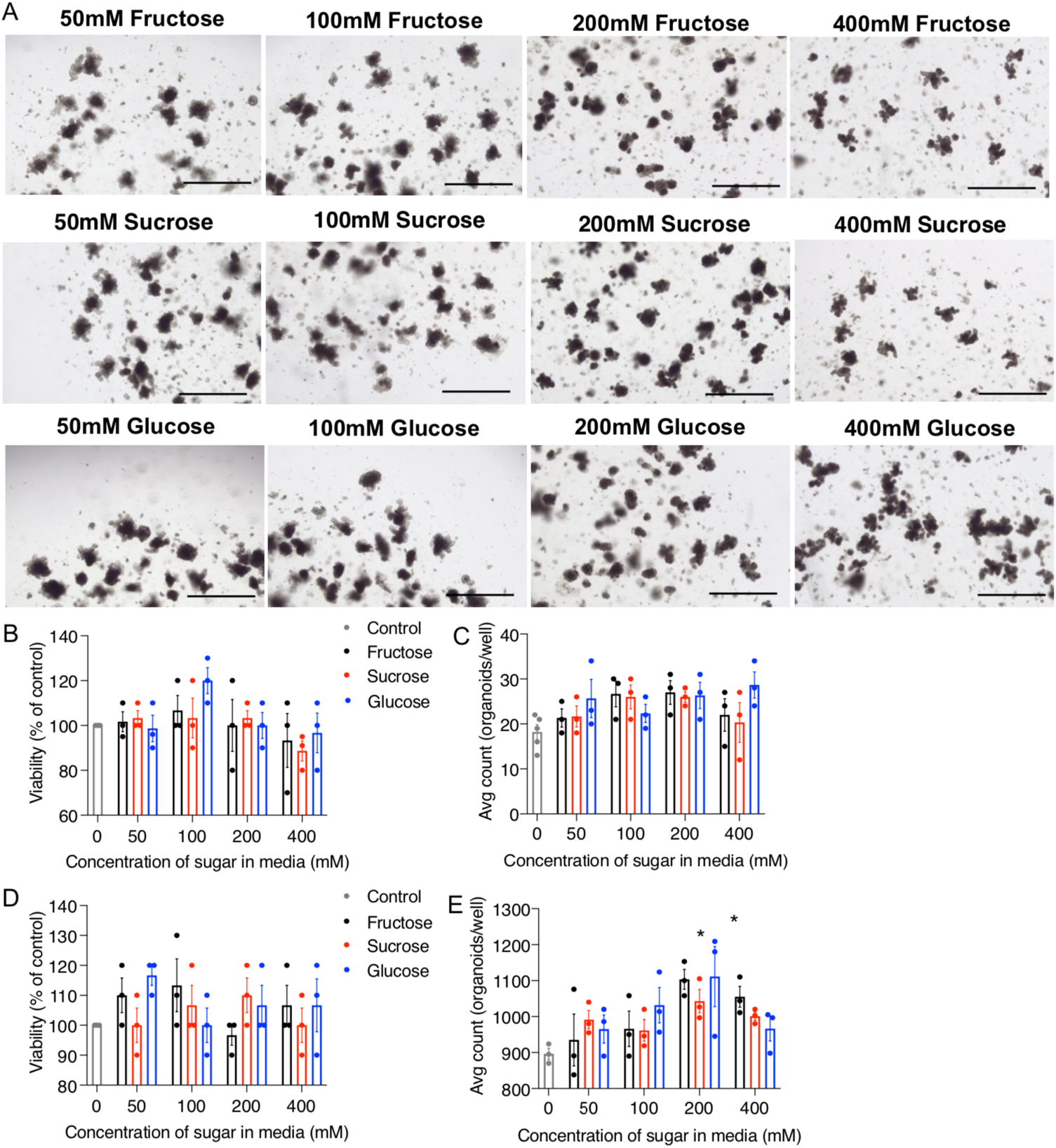
Excess sugar is not toxic to fully developed colonoids. (*A-C*) Colonic crypts were isolated from mice and cultured for 5 days into fully developed 3-D colonoids, which were then exposed to increased concentrations of sucrose for 2 days. *(A*) Representative images, (*B*) viability (percent CTG luminescence of control) and (*C*) number of organoids per well are shown. Images were taken at 4X magnification (scale bars= 200μm). (*D-E*) After developing into mature human colonoids for 5 days, excess sugar was added for 2 days. (*D*) Average viability (percent CTG luminescence of control) and (*E*) number of human colonoids are shown. (*B-E*) Data are representative of two experiments (n=3) and data points are mean +/− SEM. Stats represent one-way ANOVA with multiple comparisons to Control, where \**P*<0.05.

**Fig. S4.**
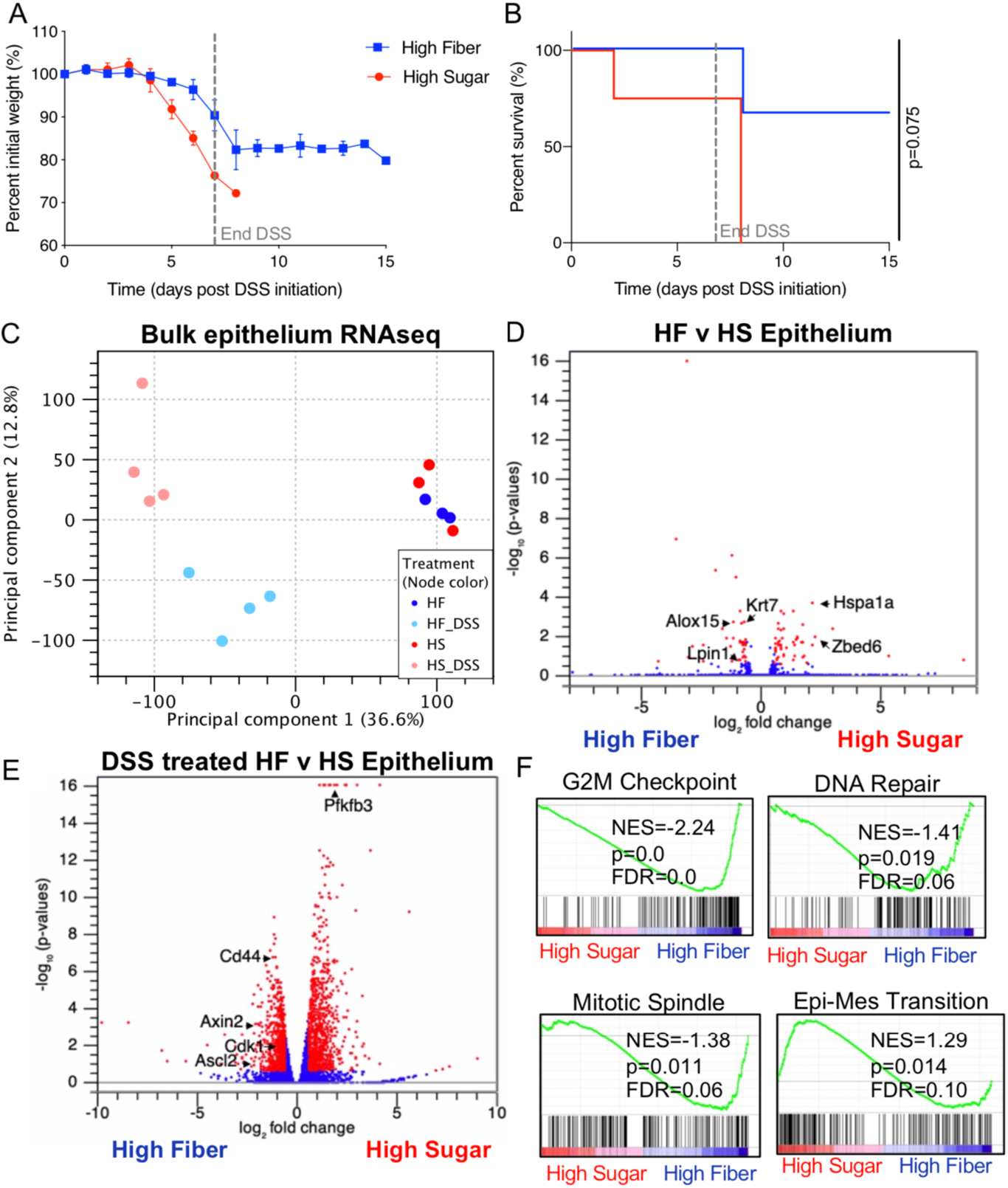
High sugar diet changes the transcriptome of the colonic epithelium. (*A-B*) *Rag1^-/-^* mice were fed HF or HS diet for 2 weeks then exposed to 3% DSS drinking water for 7 days. (*A*) Percent initial weight and (*B*) survival shown (n=3-4, mean+/− SEM). (*C-F*) Bulk colonic epithelium was isolated from *Rag1*^-/-^ female mice fed HS or HF diet for 2 weeks with or without 3 days of 3% DSS treatment and transcriptome was sequenced (n=3-4). (*C*) PCA plot showing variance, (*D*) volcano plot for genes comparing non-DSS treated samples and (*E*) volcano plot for genes comparing DSS-treated samples are shown. (*F*) Gene set enrichment analysis (GSEA) of colonic epithelium RNAseq data showing enrichment of genes in HF/DSS treated or HS/DSS treated mice for gene sets as indicated. (*C*) PCA plot showing variance across groups with percentages on axes representing percent variance explained by each principle component. (*D, F*) Volcano plots highlight significantly differentially expressed genes in red(absolute fold change greater than 1.5, *\*P<*0.05 and FDR<0.25). Specific genes are called out with arrows.

**Fig. S5.**
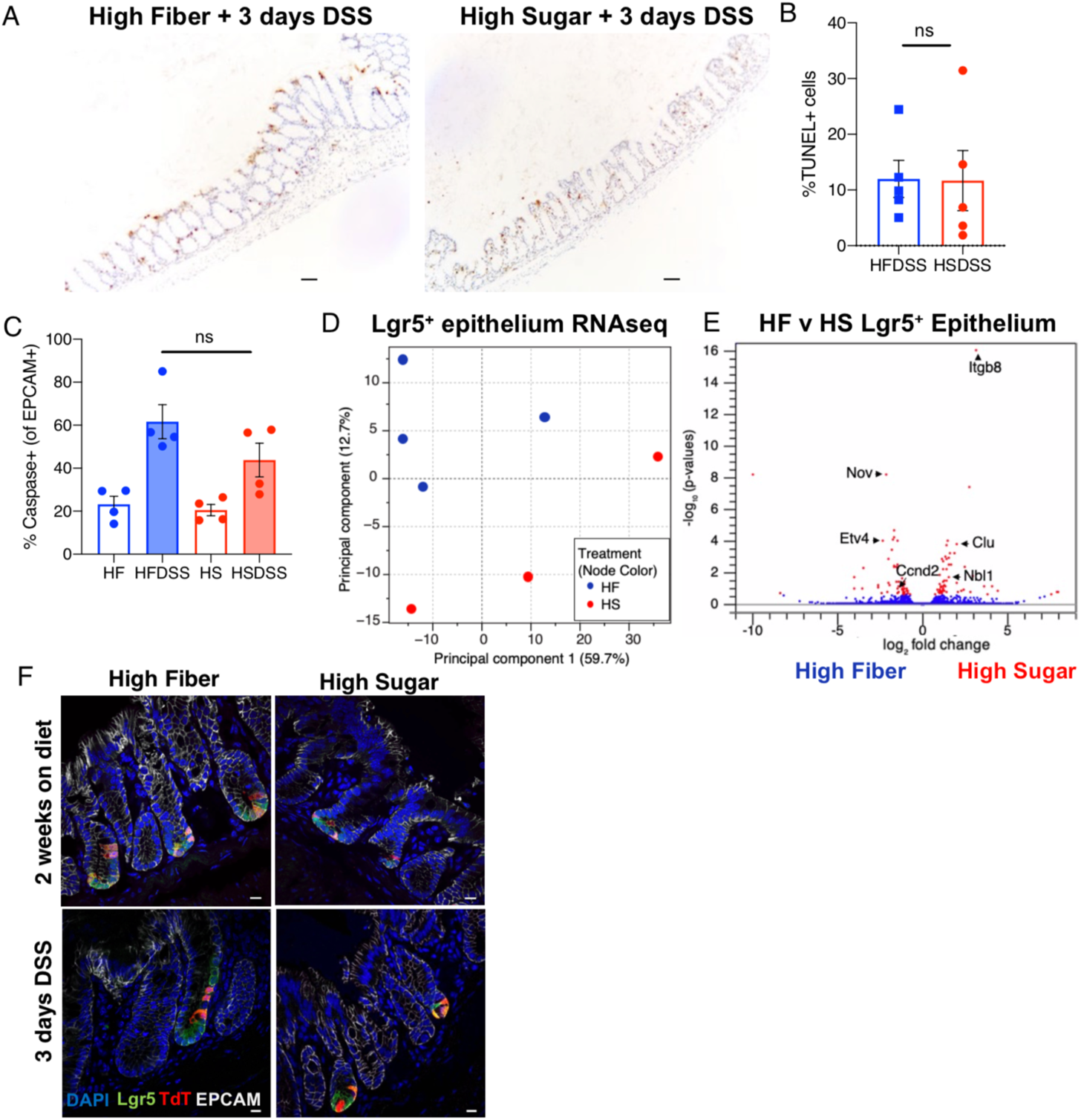
High sugar diet does not induce greater epithelial cell death, rather the transcriptome of intestinal stem cells is directly affected by excess sugar. Representative TUNEL (Terminal deoxynucleotidyl transferase dUTP nick end labeling) of colonic sections after 3 days of 3% DSS treatment in HF and HF-fed mice. Images were taken at ×10 magnification, scale bars represent 100μm. (A) Percent of cells that are TUNEL^+^. Data are representative of one experiment (n=5) and data points represent individual mouse and error bars represent SEM. (B) Mice were fed HS or HF diet and treated 3 days with 3% DSS, colonic epithelium was isolated stained with Caspase-3 for flow cytometry analysis. Data represent percent of EPCAM^+^ cells that are Caspase-3^+^ and are representative of two independent experiments (n=4). (*D-E*) Lgr5^+^ (GFP^+^) colonic epithelial cells were sorted from HS or HF-fed *Lgr5^eGFP-cre-ERT2^* reporter female mice and analyzed by RNAseq (n=3-4). (*D*) PCA plot showing variance across groups with percentages on axes representing percent variance explained by each principle component. (*E*) Volcano plots highlight significantly differentially expressed genes in blue (absolute fold change greater than 1.5, *\*P<*0.05 and FDR<0.25). Distinct genes are called out with arrows. (F) Lgr5^eGFP-Cre-ERT2^ *Rosa^LSL-TdTomato^* mice were fed HS or HF diet for 2 weeks, injected with tamoxifen on day 1 of DSS treatment and sacrificed on day 3 of DSS for Lgr5^+^ ISC lineage tracing. Representative images of colonic crypts with Lgr5^eGFP^ (green) and Tomato^+^ progeny (red). Images were taken at 60X magnification, scale bars represent 10μm.

**Fig. S6.**
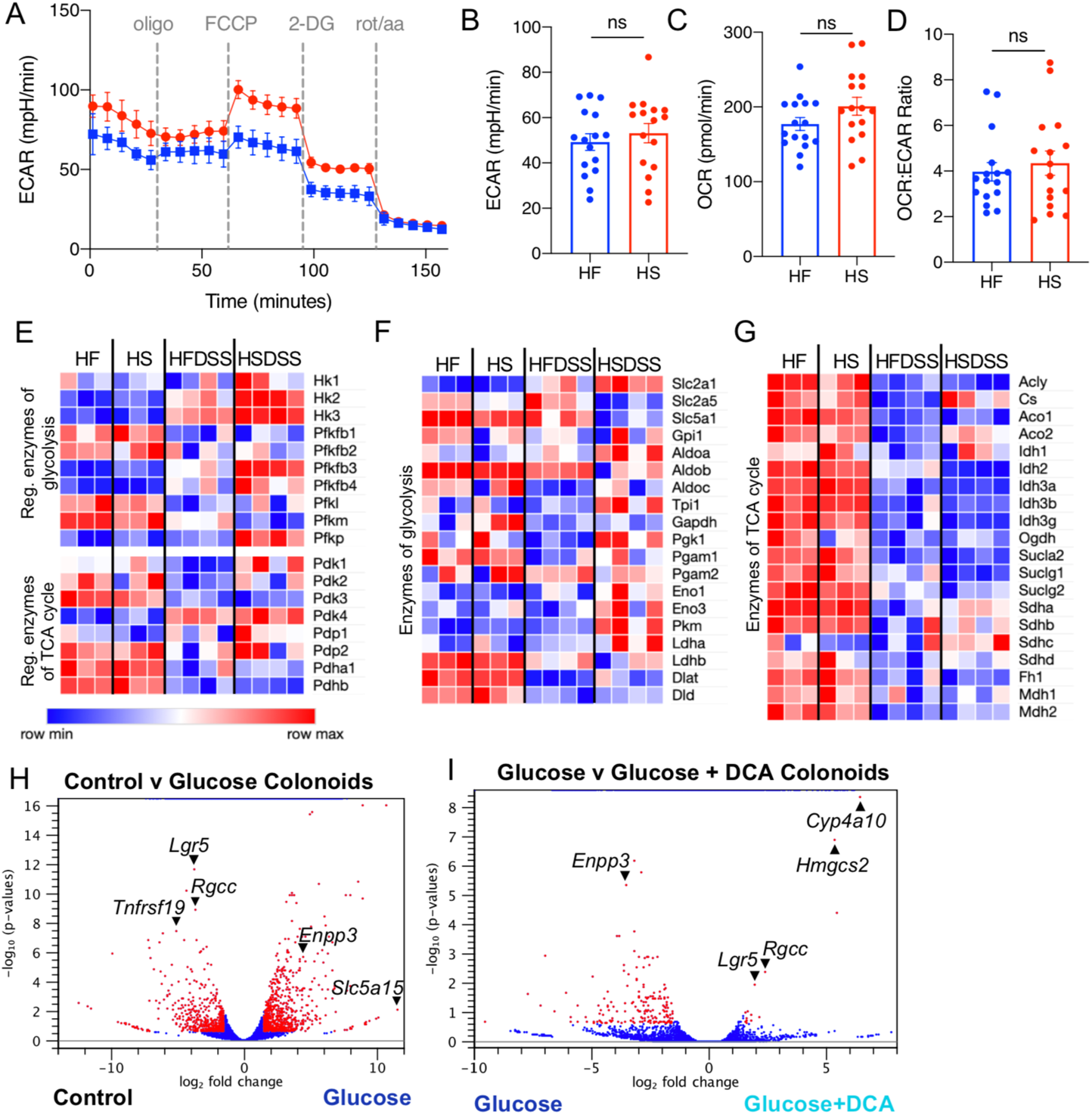
High sugar diet alters the metabolism colonic crypt epithelial cells. (*A-D*) Colonic crypts were isolated from HS or HF-fed mice. (*A*) Representative trace of extracellular acidification rate (ECAR) of crypts after 30 minutes of equilibration. (*B-C*) Tabulated data represents basal OCR and ECAR after 30 minutes equilibration (from OCR trace in Fig. 4*A*). (*D*) OCR to ECAR ratio, using basal rates from (*B*) and (*C*). (*E-G*) Heatmap of (*E*) rate-limiting enzymes in glycolysis and TCA pathways, (*F*) enzymes of glycolysis and (F) TCA cycle pathway from RNAseq of colonic epithelium from HS or HF-fed mice, with and without 3 days of 3% DSS treatment, where red and blue represent high or low expression level, respectively, normalized across rows. (*H-I*) Isolated colonic crypts were cultured in no-added sugar (Control) or 70mM of glucose (Glucose), with or without DCA (dichloroacetate) for 4 days. Colonoids were isolated in Trizol and analyzed via RNAseq. Volcano plots highlight significantly differentially expressed genes in red (absolute fold change greater than 1.5, p<0.05 and FDR<0.25) for (*H*) Control versus Glucose treated and (*I*) Glucose versus Glucose/DCA treated colonoids. Distinct genes are called out with arrows.

